# Functional analyses of phosphatidylserine/PI(4)P exchangers with diverse lipid species and membrane contexts set unanticipated rules on lipid transfer

**DOI:** 10.1101/2021.07.16.452025

**Authors:** Souade Ikhlef, Nicolas-Frédéric Lipp, Vanessa Delfosse, Nicolas Fuggetta, William Bourguet, Maud Magdeleine, Guillaume Drin

## Abstract

Several members of the oxysterol-binding protein-related proteins (ORPs)/oxysterol-binding homology (Osh) family exchange phosphatidylserine (PS) and phosphatidylinositol 4-phosphate (PI(4)P) at the endoplasmic reticulum/plasma membrane (PM) interface. It is unclear whether they preferentially exchange PS and PI(4)P with specific acyl chains to tune the features of the PM, whether they use phosphatidylinositol 4,5-bisphosphate (PI(4,5)P_2_) instead of PI(4)P for exchange processes and whether their activity is influenced by the association of PS with sterol in the PM. Here, we measured *in vitro* how the yeast Osh6p and human ORP8 transported diverse PS and PI(4)P subspecies, including major cellular species, between membranes. We established how their activity is impacted by the length and unsaturation degree of these lipids. Surprisingly, the speed at which they individually transfer these ligands inversely depends on their affinity for them. To be fast, the transfer of high-affinity ligands requires their exchange for a counterligand of equivalent affinity. Besides, we determined that Osh6p and ORP8 cannot use PI(4,5)P_2_ for intracellular lipid exchange, as they have a low affinity for this lipid compared to PI(4)P, and do not transfer more PS into sterol-rich membranes. This study provides insights into PS/PI(4)P exchangers and sets unanticipated rules on how the activity of lipid transfer proteins relates to their affinity for ligands.

## Introduction

Lipid transfer proteins (LTPs) are cytosolic proteins that distribute diverse lipids between organelles, and along with metabolic pathways, regulate the features of cell membranes^1–10^. Some members of a major family of LTPs, the oxysterol-binding protein-related proteins (ORP)/oxysterol-binding homology (Osh) family, vectorially transfer lipids by exchange mechanisms^11^. In yeast, Osh6p and its closest homologue Osh7p transfer phosphatidylserine (PS), an anionic lipid made in the endoplasmic reticulum (ER), to the plasma membrane(PM)^12^, where this lipid must be abundant to support signaling pathways^13^.

Crystallographic data has revealed that Osh6p consists of one domain – called ORD (OSBP-related domain) – with a pocket that could alternately host one molecule of PS or PI(4)P, a lipid belonging to the class of phosphorylated phosphoinositides (PIPs) ^12,13^. The pocket is closed by a molecular lid once the lipid is loaded. These structural data along with *in vitro* analyses and cellular observations have revealed the following mechanism: Osh6/7p extract PS from the ER and exchange it for PI(4)P at the PM; then they deliver PI(4)P into the ER and take PS once again. This PS/PI(4)P exchange cycle is propelled by the synthesis of PI(4)P from phosphatidylinositol (PI) in the PM and its hydrolysis in the ER membrane, which maintains a PI(4)P concentration gradient between the two compartments.

The PS/PI(4)P exchange mechanism is evolutionarily conserved^14^. Human cells express ORP5 and ORP8 that tether the ER membrane to the PM and exchange PS and PI(4)P between these membranes. They include an N-terminal pleckstrin homology (PH) domain, an ORD resembling Osh6p, and a C-terminal transmembrane segment^15,16^. They are anchored to the ER by this segment and associate with the PM via their PH domain that targets PI(4)P but also PI(4,5)P_2_ ^17–19^, which is another essential PIP of the PM^20,21^. Recently, a complex relationship has been unveiled between the ORP/Osh-mediated PS transfer process and the PI(4,5)P_2_-dependent signaling competences of the cell ^17,19,22^. It lies in the fact that PI(4)P is both used as a precursor for PI(4,5)P_2_ production and a fuel for PS/PI(4)P exchange. Moreover, as PI(4,5)P_2_, like PI(4)P, serves as anchoring point at the PM for ORP5/8, its level in the PM controls the recruitment and therefore the exchange activity of these LTPs^14,19^. Consequently, PS/PI(4)P exchange allows for a reciprocal control of PS delivery and PI(4,5)P_2_ synthesis in the PM^19,22^.

Several functional aspects of PS/PI(4)P exchangers remain enigmatic. First, it is unclear whether they selectively transfer certain PS and PI(4)P subspecies at the ER/PM interface. Eukaryotic cells contain a repertoire of PS and PIP subspecies with acyl chains of different lengths and unsaturation degrees. The nature and proportion of each subspecies in these repertoires vary considerably between organisms (*e.g.* yeasts and mammals^23–25^) but also cell types and tissues in mammals^26,27^. Moreover, inside the cell, the relative proportion of each PI(4)P and PS subspecies differs among organelles^28,29^. Some of these subspecies are predominant and this might have functional reasons. For instance, certain unsaturated PS species seem to associate preferentially with sterol in the PM, which could control the transversal distribution of sterol and the lateral distribution of PS ^30,31^, and thereby the asymmetry and remodeling propensity of this membrane. Also functional lipid nanodomains are formed by the association of 18:0/18:1-PS with very-long-chain sphingolipids^25^. A hallmark of mammal cells is the dominance of polyunsaturated PIPs with 18:0/20:4 acyl chains. This seems critical for the maintenance of a PI(4,5)P_2_ pool and a PI(4,5)P_2_-dependent signaling process in the PM via the so-called PIP cycle^32^. Consequently, it is worth analyzing how ORP/Osh proteins transfer PS and PI(4)P species with different acyl chains in order to define to what extent they can contribute to the tuning of lipid homeostasis in the PM.

A second issue concerns the links between the ORP/Osh-mediated PS transfer process and the regulation of PI(4,5)P_2_ levels. It has been reported recently that ORP5/8 use PI(4,5)P_2_ rather than PI(4)P as a counterligand for supplying the PM with PS^17^. This would mean that the PI(4,5)P_2_ level in the PM is directly lowered by the consumption of PI(4,5)P_2_ during exchange cycles. Yet this conclusion is disputed and remains surprising in view of the very first structural analyses that suggest that the polar head of PI(4,5)P_2_, unlike that of PI(4)P, cannot be accommodated by the ORD due to steric constraints^33^. The structures of the ORD of ORP1 and ORP2 in complex with PI(4,5)P_2_ have been solved^34,35^ but they revealed that the PI(4,5)P_2_ molecule is only partially buried in the binding pocket. All these observations raise doubts about the existence of functional PI(4,5)P_2_-bound forms of ORPs, including ORP5 and ORP8, in the cell.

Third, as mentioned above, unsaturated PS and sterol preferentially associate with each other in the PM^30,31^. Recently, it has been proposed that unsaturated PS and PI(4)P co-distribute into sterol-rich areas in the PM of yeast to form nanodomains that are preferentially targeted by PI(4)P 5-kinase and are the place of active PI(4)P-to-PI(4,5)P_2_ conversion^22^. Osh proteins, such as Osh6/7p but also Osh4/5p which are sterol/PI(4)P exchangers, control the formation of these domains. One might wonder whether the tight association of sterol with PS acts as a thermodynamic trap that aids PS/PI(4)P exchangers to accumulate PS in the PM and thus contribute to the coupling between PS transfer and PI(4,5)P_2_ synthesis.

Here, we addressed these three interrelated questions by conducting *in vitro* functional analyses of Osh6p and ORP8, combined with simulations and cellular observations. Using a large set of PS subspecies, we found that these LTPs transfer unsaturated PS more slowly than saturated PS between liposomes. In contrast, in a situation of PS/PI(4)P exchange, only the transfer of unsaturated PS species is largely accelerated, and efficient exchange occurs with certain unsaturated PS and PI(4)P species that are prominent in cells. Unexpectedly, by measuring the affinity of Osh6p and ORP8 for PS and PI(4)P subspecies and correlating these data with transfer rates, we established that the simple transfer of high-affinity ligands is slower than that of low-affinity ligands. Next we found that high-affinity ligands are rapidly transferred only if they can be exchanged for ligands of equivalent affinity. Furthermore, we determined that, if PI(4)P and PI(4,5)P_2_ are both accessible to Osh6p and ORP8, PI(4,5)P_2_ cannot be transferred or exchanged for PS because PI(4,5)P_2_ is a low-affinity ligand. This suggests that PI(4,5)P_2_ cannot be transported by ORP/Osh proteins in cells. Finally, we found that the activity of PS/PI(4)P exchangers barely changes on sterol-rich membranes. Our study provides insights into PS/PI(4)P exchangers but also sets general rules on how the activity of LTPs relates to their affinity for lipids, which improves our knowledge of lipid transfer.

## Results

### Osh6p and ORP8 transfer saturated and unsaturated PS species differently

We first measured *in vitro* the speed at which Osh6p and the ORD of ORP8 (ORP8[370-809]^14^, hereafter called ORD8) transferred PS subspecies with different acyl chains between two membranes (**Figure 1a**). Our series comprised subspecies with saturated acyl chains of increasing length (12:0/12:0, 14:0/14:0, 16:0/16:0, 18:0/18:0), with two C18 acyl chains that are more or less unsaturated (18:0/18:1, 18:1/18:1, 18:2/18:2) and with one saturated C16 acyl chain at the sn-1 position and one C18 acyl chain, with a different unsaturation degree, at the sn-2 position (16:0/18:1, 16:0/18:2). Note that 16:0/18:1-PS is the dominant PS species in *S. cerevisiae* yeast^22,24,28,36^ (under standard growing conditions) whereas in humans, 18:0/18:1-PS and 16:0/18:1-PS are the two most abundant species^25,37–39^. PS transfer was measured between L_A_ liposomes, made of DOPC and containing 5% PS (mol/mol) and 2% Rhod-PE, and L_B_ liposomes only made of DOPC, using the fluorescent sensor NBD-C2_Lact_. In each measurement, NBD-C2_Lact_ was initially bound to L_A_ liposomes and its fluorescence was quenched due to energy transfer to Rhod-PE; when LTP transferred PS to L_B_ liposomes, NBD-C2_Lact_ translocated onto these liposomes and the fluorescence increased **(Figure 1b)**. By normalizing the NBD signal, we established transfer kinetics **(Figure 1 – Figure Supplement 1a)** and initial transfer rates **(Figure 1c)**. Osh6p transferred saturated PS rapidly, with rates between 7.6 ± 1 (mean ± s.e.m.,14:0/14:0-PS) and 35.2 ± 2 PS.min^−1^ (18:0/18:0-PS). The transfer of unsaturated PS species was much slower (from 0.8 ± 0.1 to 3.1 ± 0.8 PS.min^−1^ *per* Osh6p). A different picture was obtained when we measured PS transfer in a situation of PS/PI(4)P exchange using L_B_ liposomes containing 5% 16:0/16:0-PI(4)P. The transfer rates measured with 14:0/14:0-PS and 16:0/16:0-PS were similar to those measured in non-exchange contexts, and significantly lower with 12:0/12:0-PS and 18:0/18:0-PS. In contrast, the transfer rate of unsaturated PS, with the exception of 18:2/18:2-PS, strongly increased (by a factor from 3.2 to 19.5).

**Figure 1.**
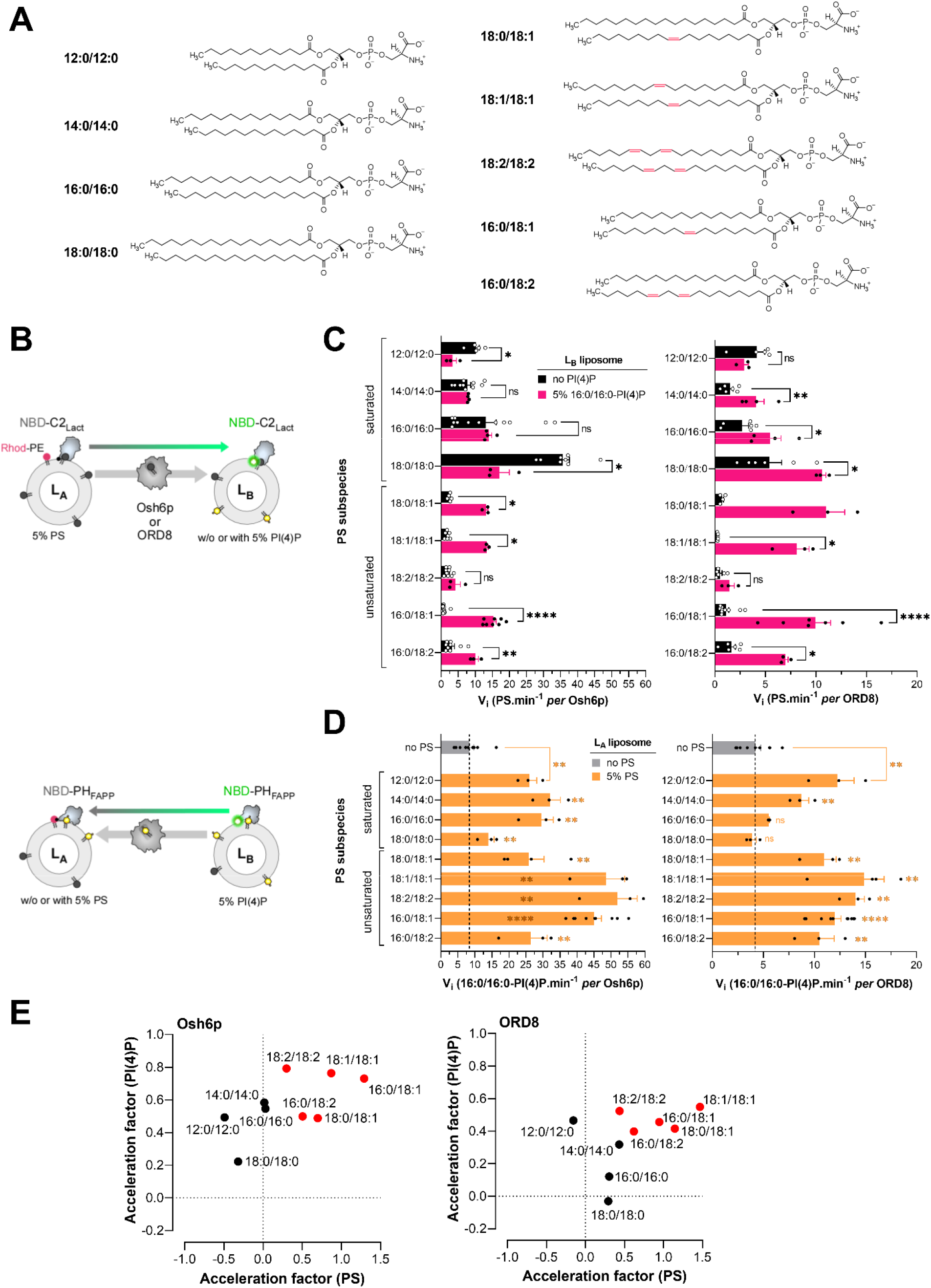
PS transfer and PS/PI(4)P exchange activity of Osh6p and ORD8 measured with different PS subspecies. **(A)** Name and chemical structure of the different PS subspecies. **(B)** Description of FRET-based protocols to measure PS transfer from L_A_ to L_B_ liposomes using the NBD-C2_Lact_ sensor, and PI(4)P transfer along the opposite direction using NBD-PH_FAPP_. **(C)** Initial transfer rates determined for each PS subspecies. Osh6p (200 nM) or ORD8 (240 nM) was added to L_A_ liposomes (200 µM total lipid), made of DOPC and containing a given PS species at 5%, mixed with L_B_ liposomes (200 µM) containing or not 5% 16:0/16:0-PI(4)P and 250 nM NBD-C2_Lact_. Data are represented as mean ± s.e.m. (Osh6p, non-exchange condition, n = 3-16; Osh6p, exchange condition, n = 3-8; ORD8, non-exchange condition, n = 3-11; ORD8, exchange condition, n = 3-7). Statistically significant differences between PS transfer rates measured under non-exchange and exchange conditions were determined using an unpaired Mann–Whitney U test; **** p<0.001, \*\**p* < 0.01, \**p* < 0.05, ns: not significant. **(D)** Initial PI(4)P transfer rate. L_B_ liposomes containing 5% 16:0/16:0-PI(4)P were mixed with L_A_ liposomes, containing or not a given PS species (at 5%), and with Osh6p (200 nM) or ORD8 (240 nM) in the presence of 250 nM NBD-PH_FAPP_. Data are represented as mean ± s.e.m. (Osh6p, non-exchange condition, n = 9; Osh6p, exchange condition, n = 3-9; ORD8, non-exchange condition, n = 9; ORD8, exchange condition, n = 3-10). An unpaired Mann–Whitney U test was used to determine the statistically significant differences between the PI(4)P transfer rates measured in non-exchange and exchange conditions; **** p<0.0001, \*\**p* < 0.01, \**p* < 0.05, ns: not significant. **(E)** Acceleration of PS transfer as a function of the acceleration of PI(4)P transfer determined from rates measured in non-exchange and exchange conditions with all PS subspecies.

Overall, ORD8 transported PS more slowly than Osh6p in both exchange and non-exchange situations (**Figure 1c, Figure 1 – Figure Supplement 1b**). Nevertheless, the activity of ORD8 changed depending on the PS subspecies in a manner similar to that of Osh6p, as highlighted by the correlation of transfer rates measured for the two LTPs with each PS ligand (R^2^ ∼ 0.75, **Figure 1 – Figure Supplement 1c**). Like Osh6p, ORD8 transferred saturated PS species more rapidly than unsaturated ones when PI(4)P was absent. In a situation of PS/PI(4)P-exchange, the transfer of unsaturated PS species (except for 18:2/18:2-PS) was much more rapid (up to 29-fold) whereas the transfer of saturated PS was slightly enhanced or inhibited. Collectively, these data did not point to a monotonic relationship between PS transfer rates and the length of PS acyl chains or the degree of unsaturation of these chains. However, they indicated that PS species were transported and exchanged with PI(4)P quite differently depending on whether or not they had at least one double bond.

### Coupling between the transfer rate of PS species and PI(4)P under exchange conditions

Next, we determined whether the rate of 16:0/16:0-PI(4)P transfer was different depending on the nature of the PS species under the exchange conditions. Using a fluorescent PI(4)P sensor (NBD-PH_FAPP_) and a FRET-based strategy akin to that used to measure PS transfer, we measured the speed at which PI(4)P, at 5% in L_B_ liposomes, was transported by Osh6p and ORD8 to L_A_ liposomes devoid of PS or containing a given PS species (at 5%) (**Figure 1b, Figure 1 – Figure Supplement 2**). With PS-free L_A_ liposomes, the initial PI(4)P transfer rate was 8.4 ± 1.3 PI(4)P.min^−1^ for Osh6p and 4.2 ± 0.5 PI(4)P.min^−1^ for ORD8 (**Figure 1d**). In a situation of lipid exchange, these transfer rates increased to a different degree when L_A_ liposomes contained a PS species other than 18:0/18:0-PS and, in experiments with ORD8, 16:0/16:0-PS (**Figure 1d**). For each PS species, we calculated an acceleration factor corresponding to the ratio (expressed as a log value) of the PI(4)P transfer rate, measured in the presence of this PS species, to the PI(4)P transfer rate measured in the absence of counterligand. Also, acceleration factors based on PS transfer rates reported in **Figure 1c** were determined. Then for each LTP, these two factors were plotted against each other, allowing saturated and unsaturated PS species to cluster in two groups (**Figure 1e**). With Osh6p, the group including saturated PS species was characterized by null or negative acceleration factors for PS (down to - 0.49) associated with low or moderate acceleration factors for 16:0/16:0-PI(4)P (from 0.20 to 0.58). In contrast, the group corresponding to unsaturated PS species was characterized by higher acceleration factors for both PS (from 0.30 to 1.30) and PI(4)P (from 0.49 to 0.79). With ORD8, the acceleration factors for saturated PS were negative, null or moderate (from −0.15 to 0.47) and associated with null or moderate acceleration factors for PI(4)P (from 0 to 0.48). For unsaturated PS, the acceleration factors were higher, ranging from 0.44 to 1.47 for PS and from 0.40 to 0.55 for PI(4)P. The observation of high acceleration factors for both unsaturated PS and 16:0/16:0-PI(4)P, and much lower or even negative values for saturated PS species, suggests that LTPs exchange unsaturated PS for PI(4)P much more efficiently than saturated PS.

### Exchange activity with prominent cellular PS and PI(4)P species

We next measured how Osh6p and ORD8 exchanged PS and PI(4)P species that are dominant in the yeast and/or human repertoire. With Osh6p, we tested 16:0/18:1-PS and 18:0/18:1-PS with 16:0/18:1-PI(4)P, one of the two most abundant yeast PI(4)P species^22,23^. As a comparison we tested a non-yeast species, 18:0/20:4-PI(4)P, which is the main constituent of purified brain PI(4)P. With ORD8, we tested the two same PS species with 18:1/18:1-PI(4)P that resembles unsaturated PI(4)P species (36:1 and 36:2) found in transformed cells^40,41^ and 18:0/20:4-PI(4)P, which is prominent in primary cells and tissues^41^. As a comparison we used 16:0/16:0-PI(4)P as in our previous assays.

Osh6p slowly transferred the two PS species from L_A_ to L_B_ liposomes in the absence of PI(4)P but ten times faster when L_B_ liposomes contained 16:0/18:1-PI(4)P or 16:0/16:0-PI(4)P. Smaller accelerations of PS transfer were seen with 18:0/20:4-PI(4)P as counterligand. Conversely, in the absence of PS, Osh6p hardly transported any 16:0/18:1-PI(4)P and 18:0/20:4-PI(4)P (< 0.4 lipids.min^−1^) compared to saturated PI(4)P (5.4 lipids.min^−1^) **(Figure 2a, Figure 2 – Figure Supplement 2a, b)**. When L_A_ liposomes contained PS, the transfer rate of all PI(4)P species increased but with rates that were high for 16:0/16:0-PI(4)P (25.8-39.6 lipids.min^−1^), intermediate for 16:0/18:1-PI(4)P (7-13.3 lipids.min^−1^) and low for 18:0/20:4-PI(4)P (1.63-2.4 lipids.min^−1^). Interestingly, the 16:0/18-1-PS and 16:0/18:1-PI(4)P transfer rates were both similar and high in a situation of lipid exchange, suggesting that Osh6p can execute an efficient one-for-one exchange of these major yeast PS and PI(4)P species.

**Figure 2.**
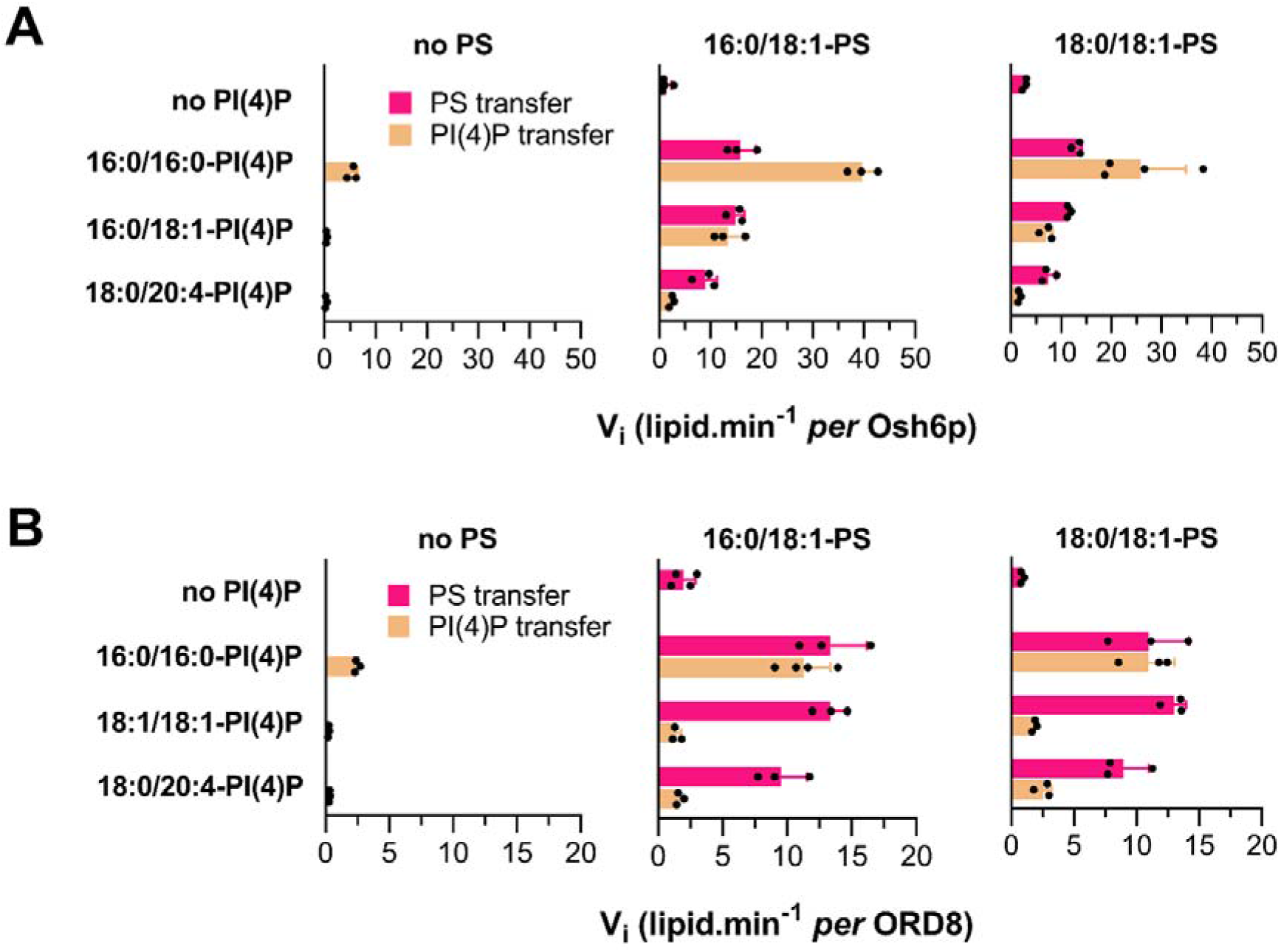
Ability of Osh6p and ORP8 to transfer and exchange cellular PS and PI(4)P species. **(A)** Initial PS and PI(4)P transfer rates, measured along opposite directions, between L_A_ and L_B_ liposomes (200 µM total lipid each) with Osh6p (200 nM) at 30°C, in the absence of counterligand or in a situation of lipid exchange, with diverse combinations of PS (16:0/18:1 or 18:0/18:1, 5% in L_A_ liposomes) and PI(4)P species (16:0/16:0, 16:0/18:1 or 18:0/20:4, 5% in L_B_ liposomes). Data are represented as mean ± s.e.m. (n = 3-4). **(B)** Similar experiments were conducted with ORD8 (240 nM) at 37°C using 18:1/18:1-PI(4)P instead of 16:0/18:1-PI(4)P. Data are represented as mean ± s.e.m. (n = 3-4).

In the absence of PI(4)P, ORD8 slowly transferred 16:0/18:1-PS and 18:0/18:1-PS between membranes and much faster in a situation of exchange, by a factor of 4.8-6.8 and 11.5-16.6, respectively, depending on the PI(4)P species used as counterligand **(Figure 2b, Figure 2 – Figure Supplement 2c, d)**. Like Osh6p, ORD8 barely transferred unsaturated PI(4)P under non-exchange conditions (< 0.26 lipids.min^−1^), compared to 16:0/16:0-PI(4)P. Under exchange conditions, ORD8 transferred these PI(4)P species more rapidly but far less than PS in the opposite direction. This suggests that ORP8 cannot efficiently exchange unsaturated PS for PI(4)P. We conclude that the acyl chain composition of PI(4)P, and primarily its unsaturation degree, impacts how Osh6p and ORP8 transfer and use this PIP in exchange for PS.

### Osh6p and ORP8 activities drastically change if the sn-1 or sn-2 chain of PS is monounsaturated

Striking differences were seen between 18:0/18:0-PS and unsaturated forms of this lipid in our transfer assays. In particular, data obtained with 18:0/18:0-PS and 18:0/18:1 PS suggested that only one double bond in PS was sufficient to drastically change LTP activity. Whether this depends on the location of this double bond in the sn-2 chain was unclear. Therefore, we compared how Osh6p transferred 18:0/18:1 PS and 18:1/18:0-PS, in which the saturated and monounsaturated acyl chains are permuted, between membranes **(Figure 3a)**. In mere transfer assays, 18:0/18:1-PS and 18:1/18:0-PS were transported at rates that were slightly different (4.7 vs 2.1 PS.min^−1^) but ten-fold more slowly than 18:0/18:0-PS **(Figure 3b)**. In the presence of 16:0/18:1-PI(4)P, the transfer of the two unsaturated PS species was faster whereas the transfer of 18:0/18:0-PS was inhibited. The opposite transfer of 16:0/18:1-PI(4)P was inhibited if 18:0/18:0-PS was tested as counterligand but enhanced using the two unsaturated PS forms **(Figure 3a)**. Similar results were obtained with ORD8 except that the transfer of 18:0/18:0-PS was slightly more rapid (by 2.2-fold) in exchange conditions (here 18:1/18:1-PI(4)P was used as counterligand **(Figure 3c))**. However, the rate of acceleration was low compared to that measured with 18:0/18:1-PS and 18:1/18:0-PS (by 24.0 and 8.4-fold, respectively). Jointly these results indicate that only one double bond, in one or the other acyl chain of PS, is enough to dramatically change how the LTPs transports and exchanges this lipid for PI(4)P.

**Figure 3.**
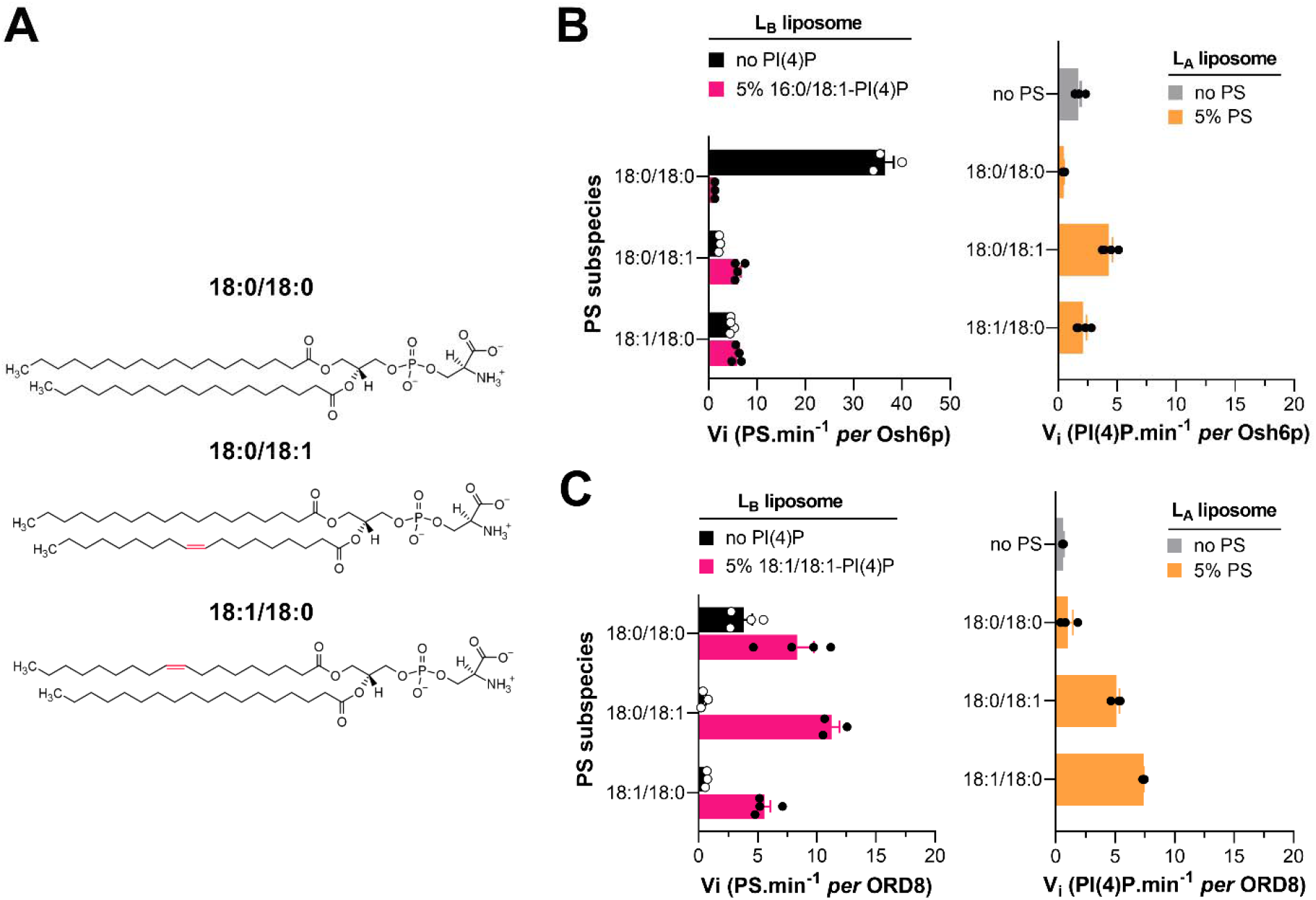
Transfer of 18:1/18:0-PS vs 18:0/18:1-PS by Osh6p and ORD8. **(A)** Chemical structure of 18:1/18:0-PS compared to that of 18:0/18:0-PS and 18:0/18:1-PS. **(B)** Initial rates of PS and PI(4)P transfer, measured along opposite directions, between L_A_ and L_B_ liposomes (200 µM total lipid each) with Osh6p (200 nM) at 30°C, in non-exchange or exchange contexts with 18:0/18:0-PS, 18:0/18:1-PS or 18:1/18:0-PS (5% in L_A_ liposomes) and 16:0/18:1-PI(4)P (5% in L_B_ liposomes). Data are represented as mean ± s.e.m. (n = 3-4). **(C)** Similar experiments were performed with ORD8 (240 nM) at 37°C using 18:1/18:1-PI(4)P instead of 16:0/18:1-PI(4)P.

### Osh6p and ORD8 have a higher affinity for unsaturated than saturated PS and PI(4)P species

Our results suggest that Osh6p and ORD8 transport and exchange PS and PI(4)P at a different speed depending on the unsaturation degree of these lipids. To further analyze why, we devised assays to determine the relative affinity of these LTPs for each PS and PI(4)P species. We established that the intrinsic fluorescence of both proteins (from tryptophan, with a maximum intensity at λ = 335 and 340 nm, respectively), was quenched by ∼25% when mixed with liposomes doped with 2% NBD-PS, a PS species whose C12:0 acyl chain at the sn-2 position bears an NBD moiety **(Figure 4 – Figure Supplement 1)**. Concomitantly, a higher NBD fluorescence was measured at λ = 540 nm. Adding each LTP to liposomes doped with 2% NBD-PC provoked a slighter decrease of tryptophan fluorescence, yet similar to the changes recorded with pure PC liposomes, and no change in NBD fluorescence. Likely, FRET exclusively occurs between these proteins and NBD-PS because this lipid is specifically trapped in their binding pocket and close to a number of tryptophan residues. Interestingly we found subsequently that Osh6p and ORP8 fluorescence, pre-mixed with NBD-PS-containing liposomes, increased when adding incremental amounts of liposomes containing unlabeled PS. This allowed us to measure how each PS species competes with NBD-PS for occupation of the Osh6p and ORD8 pocket, and thus determine the relative affinity of each ORD for different lipid ligands (**Figure 4a,b and Figure 4 – Figure Supplement 2a**). Remarkably, Osh6p had a very low affinity for 18:0/18:0-PS and 12:0/12:0-PS. It showed a higher affinity for 14:0/14:0-PS and 16:0/16:0-PS. Highest affinities were found with unsaturated PS and more particularly 16:0/18:1-PS, 18:1/18-1-PS and 18:0/18:1-PS. Interestingly, Osh6p had a higher affinity for 18:0/18:1-PS than its mirror 18:1/18:0-PS counterpart **(Figure 4b)**. Using this assay we also found that Osh6p had a high affinity for 16:0/18:1-PI(4)P and 18:0/20:4-PI(4)P and less for 16:0/16:0-PI(4)P **(Figure 4c)**. With ORD8, competition assays revealed that it had a much lower affinity for saturated PS than for unsaturated PS **(Figure 4 – Figure Supplement 2a**). ORD8 had a higher affinity for 18:1/18:1-PI(4)P and 18:0/20:4-PI(4)P than for 16:0/16:0-PI(4)P **(Figure 4 – Figure Supplement 2b)**. Collectively, our data indicated that Osh6p and ORD8 had a higher affinity for unsaturated than saturated lipid ligands.

**Figure 4.**
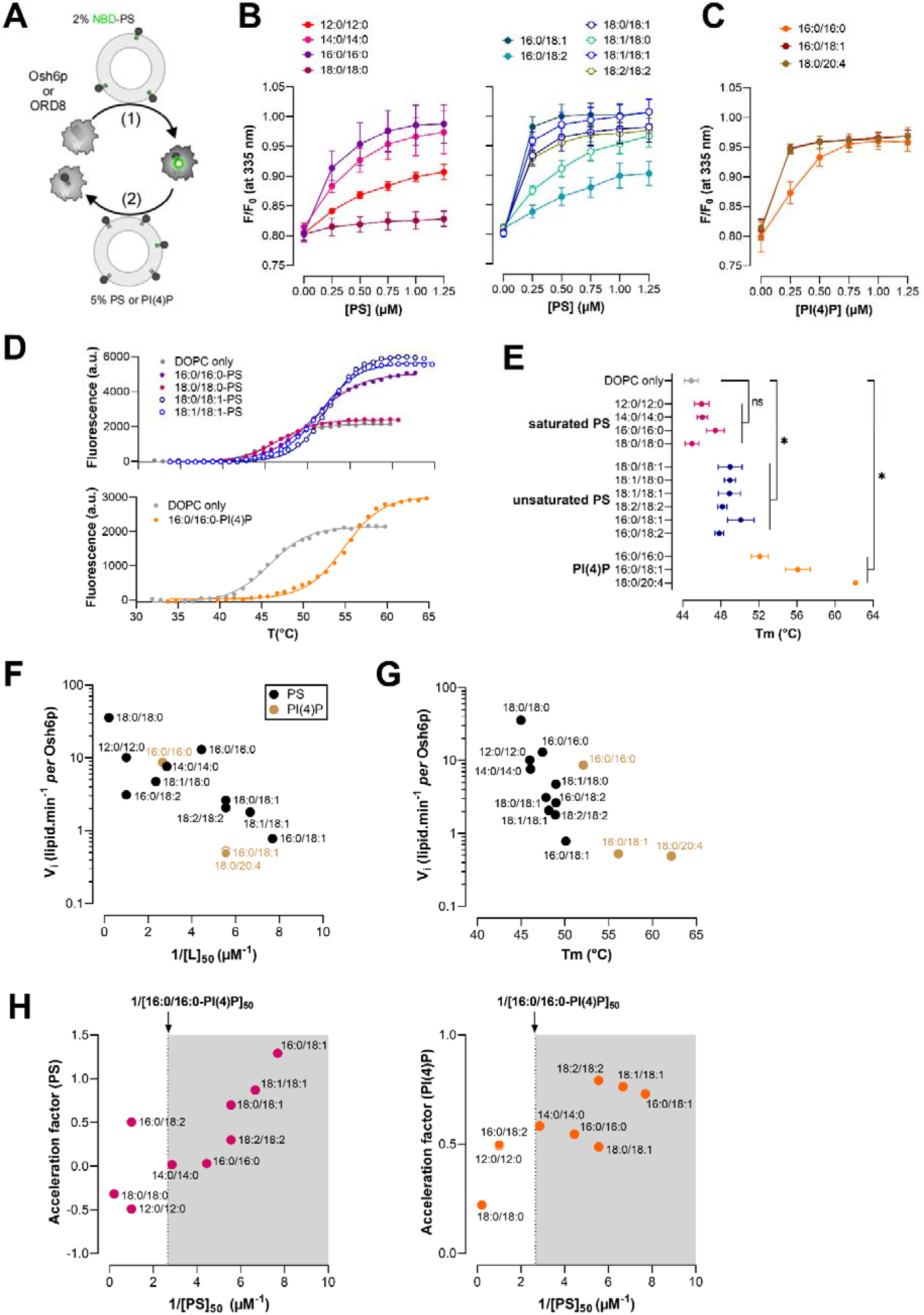
Relationship between the affinity of Osh6p for PS and PI(4)P species and its capacity to transfer them. **(A)** Principle of the NBD-PS-based competition assays. The tryptophan (W) fluorescence of Osh6p and ORD8 is quenched when they host an NBD-PS molecule. The replacement of NBD-PS by unlabeled PS restores the fluorescence of these proteins **(B)**. Competition assays with different PS species. Liposomes (100 µM total lipid, final concentration), made of DOPC and doped with 2% NBD-PS, were added to Osh6p (240 nM) in HK buffer at 30°C. The sample was excited at 280 nm and the emission was measured at 335 nm. Incremental amounts of liposome, containing a given PS species at 5%, were added to the sample. The fluorescence was normalized considering the initial F_max_ fluorescence, prior to the addition of NBD-PS-containing liposomes, and the dilution effects due to liposome addition. Data are represented as mean ± s.e.m. (n = 3). **(C)** Competition assays with liposomes containing either 5% 16:0/16:0-PI(4)P, 16:0/18:1-PI(4)P or 18:0/20:4-PI(4)P. Data are represented as mean ± s.e.m. (n = 4 for 16:0/18:1-PI(4)P, n = 3 for other PI(4)P species). **(D)** Melting curves of Osh6p loaded with different PS species or 16:0/16:0-PI(4)P. In a typical measurement, a sample containing 5 µM of protein and 5× SYPRO Orange in HK buffer was heated and fluorescence was measured at λ_em_ = 568 nm (λ_ex_ = 545 nm). A control experiment with Osh6p incubated with DOPC liposomes devoid of lipid ligands is shown (DOPC only). Only a few curves corresponding to representative ligands are shown for clarity.**(E)** Melting temperatures□(T_m_) determined for Osh6p pre-incubated with pure DOPC liposome or loaded with diverse PS and PI(4)P subspecies. Data are represented as mean ± s.e.m (n = 3-5). Pairwise comparison by unpaired Mann-Whitney U test of T_m_ rate measured with Osh6p in apo form (DOPC only) and Osh6p loaded with saturated or unsaturated PS species, or a PI(4)P species; \**p* < 0.05, ns: not significant. **(F)** Initial transfer rates determined in non-exchange contexts for PS and PI(4)P subspecies with Osh6p (shown in Figure. 1c, 2 and 3) as a function of 1/[L]_50_ values determined for each lipid subspecies. **(G)** PS transfer rates in non-exchange conditions as a function of Tm values determined with Osh6p. **(H)** Acceleration factors determined for PS and PI(4)P from experiments shown in Figure 1b and c, as a function of the 1/[L]_50_ values determined for each PS subspecies. The 1/[L]_50_ value determined with 16:0/16:0-PI(4)P is indicated.

Alternatively, Osh6p was incubated with liposomes doped with a given PS or PI(4)P subspecies, isolated and then subjected to thermal shift assays (TSAs) to evaluate to what extent it formed a stable complex with each ligand **(Figure 4d,e)**. Low melting temperatures (T_m_) were observed with Osh6p exposed to liposomes containing saturated PS species (from 45 to 47.4°C), near the T_m_ value of Osh6p incubated with DOPC liposomes devoid of ligand (44.9 ± 0.7°C). In contrast, significantly higher values were obtained with unsaturated PS (from 47.8 to 50.1°C). The highest T_m_ values were found with Osh6p loaded with 16:0/18:1 and 18:0/20:4-PI(4)P species (56.1 and 62.1°C, respectively), and a slightly lower T_m_ was found with 16:0/16:0-PI(4)P (52°C). These results suggest that Osh6p is more prone to capture and hold unsaturated PS and PI(4)P than saturated PS, corroborating the results from competition assays.

### The affinity of Osh6p and ORP8 for PS and PI(4)P species dictates how they transfer and exchange them

Next, we analyzed how the affinity of Osh6p and ORD8 for lipid ligands was related to their capacity to transfer them. To do so, we plotted the PS and PI(4)P transfer rates measured in non-exchange contexts (reported in **Figure 1c,d, Figure 2 and Figure 3**) as a function of 1/[L]_50_, with [L]_50_ being the concentration of each species necessary to displace 50% NBD-PS from each LTP in the competition assays. Remarkably, this revealed an inverse relationship between the transfer rates and 1/[L]_50_ values (**Figure 4f, Figure 4 – Figure Supplement 2c**) for each LTP. For Osh6p, plotting the transfer rates as a function of Tm values uncovered a comparable relationship (**Figure 4g)**. This suggested that the less affinity these LTPs have for a ligand, the faster they transfer it between membranes. We also plotted the acceleration factors established under exchange conditions with different PS species and 16:0/16:0-PI(4)P (showed in **Figure 1e**) against 1/[L]_50_ values determined with each PS species and LTP (**Figure 4h, Figure 4 – Figure Supplement 2d**). Interestingly a positive relationship was found between the 1/[L]_50_ values and each of these factors. Moreover, we noted that PS and PI(4)P transfer rates measured under exchange conditions were overall higher when the LTPs had a higher affinity for PS than PI(4)P (1/[PS]_50_ > 1/[16:0/16:0-PI(4)P]_50_). Collectively, these analyses reveal an inverse correlation between the affinity of an LTP for a ligand and its ability to simply transfer it down its concentration gradient. Moreover they indicate that in a situation of lipid exchange, the acceleration of the PS and PI(4)P transfer rate is more pronounced with high-affinity PS species.

### Simulation of transfer and exchange activity of the ORD as a function of its affinity for PS and PI(4)P

To understand why the affinity of Osh6p and ORD8 for PS and PI(4)P species governed how they transferred and exchanged these lipids, we built a simplified kinetic model (**Figure 5a**). It was assumed that the ORD interacts similarly with A and B membranes during a transfer process (k_ON-Mb=_ 10 s^−1^and k_OFF-Mb_=0.1 s^−1^) with an equal ability to capture and release a given lipid (similar k_ON-lipid_ and k_OFF-lipid_). We simulated initial PS transfer rates for k_ON-PS_ values ranging from 10^−2^ to 10^4^ µM^−1^.s^−1^ to evaluate how the affinity of the ORD for PS (proportional to k_ON-PS_) governs how it transfers this lipid. The k_ON-PI4P_ values was set to 10 µM^−1^.s^−1^ and k_OFF-PS_ and k_OFF-PI4P_ values to 1 s^−1^; the ORD concentration was 200 nM, with an A membrane including 5 µM accessible PS and a B membrane devoid of ligand (as in our transfer assays). A bell-shaped curve (**Figure 5b**, left panel, black dots) was obtained with a maximum at k_ON-PS_ = 3.7 µM^−1^.s^−1^ and minima near zero for very low and high k_ON-PS_ values. Remarkably our simulations indicate that an LTP can transfer a low-affinity ligand more rapidly than a high-affinity one, as seen for instance when comparing rates at k_ON-PS_ = 10 and 100 µM^−1^.s^−1^, and as observed experimentally with saturated and unsaturated PS.

**Figure 5.**
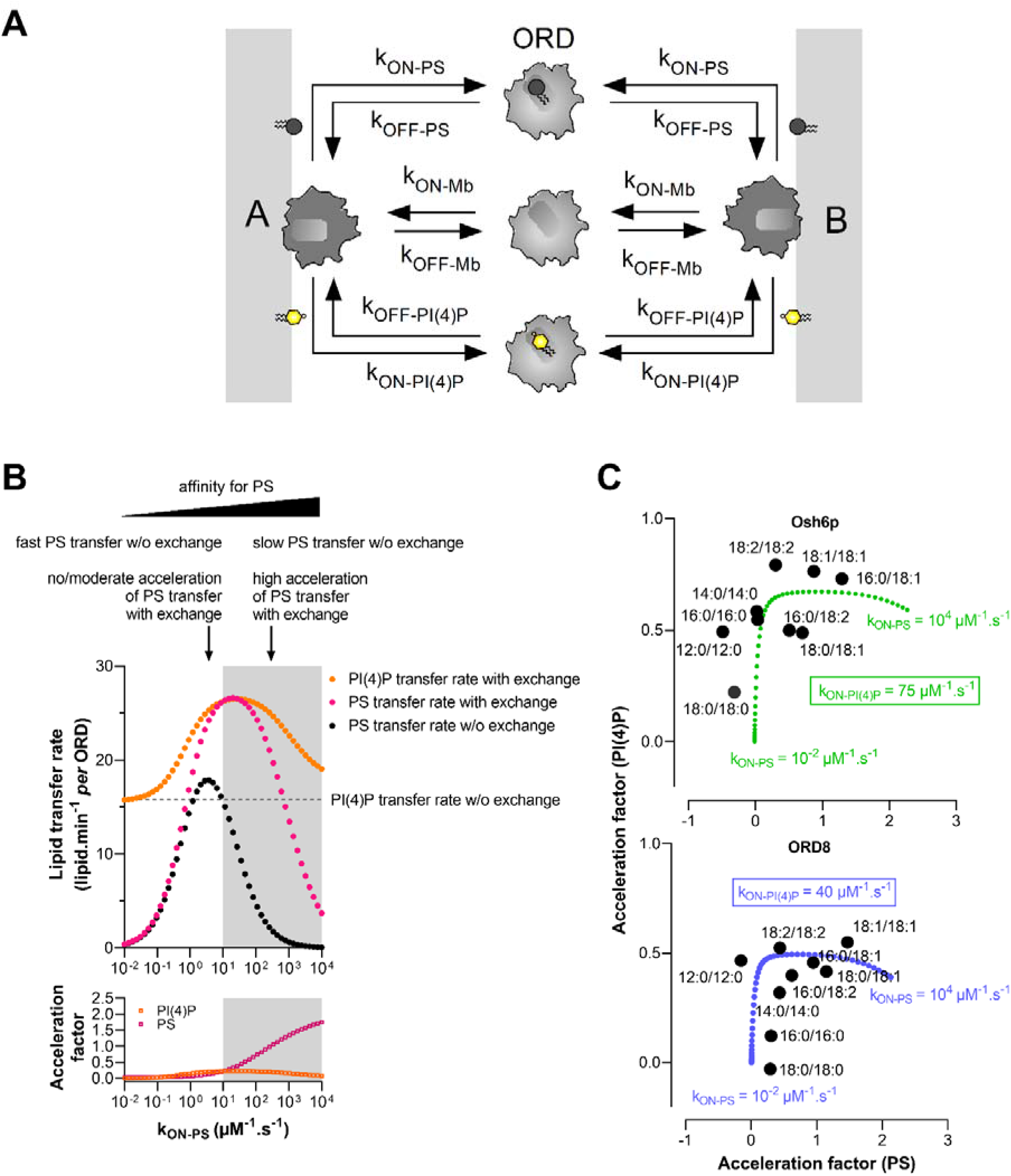
Analysis of the relation between ORD’s affinity for PS and PI(4)P and its ability to transfer these lipids between membranes. **(A)** Description of the kinetic model. Osh6p or ORD8 (ORD) interacts with the same affinity with two distinct membranes A and B, each harboring a PS and PI(4)P pool, and can extract and release PS and PI(4)P. ORD-PS and ORD-PI(4)P correspond to the ORD in 1:1 complex with PS and PI(4)P, respectively. All k_ON_ and k_OFF_ rates were set to 10 µM^−1^.s^−1^, and 1 s^−1^, respectively, unless otherwise specified. **(B)** Initial PS transfer rate (grey dots) as a function of k_ON-PS_ values (ranging from 0.01 to 10,000 µM^−1^.s^−1^), under the condition where the A membrane initially contained 5% PS and B membrane was devoid of PI(4)P (non-exchange condition). Initial PS (pink dots) and PI(4)P transfer rates (orange dots) were also calculated as a function of k_ON-PS_, considering that PS and PI(4)P were initially present at 5% in the A and B membranes, respectively (exchange condition). PI(4)P transfer rate simulated with A membrane devoid of PS (non-exchange condition) was indicated by a dashed line. The grey areas correspond to regimes where k_ON-PS_ > k_ON-PI(4)P_ *i.e.,* the ORD has more affinity for PS than PI(4)P. The acceleration factors, calculated for PS and PI(4)P, correspond to the ratio (in log value) between the transfer rates derived from simulations performed in exchange and non-exchange conditions. **(C)** Acceleration factors of PS and PI(4)P transfer in exchange conditions, established for Osh6p and ORD8 with different PS species and 16:0/16:0-PI(4)P, are plotted against each other as in the Figure 1E. For comparison, theoretical acceleration factors of PS and PI(4)P transfer, considering k_ON-PS_ value ranging from 0.01 to 10,000 µM^−1^.s^−1^ and a k_ON_-_PI(4)P_ value of 75 µM^−1^.s^−1^ for Osh6p or 40 µM^−1^.s^−1^ for ORD8, are plotted against each other (Osh6p, green; ORD8, blue).

Next, using the same range of k_ON-PS_ values, we simulated PS and PI(4)P transfer rates in a situation of lipid exchange between A and B membranes that initially contained 5 µM PS and PI(4)P, respectively (**Figure 5b**, left panel, pink dots for PS and orange dots for PI(4)P). Acceleration factors were determined from rates established for each lipid in exchange and non-exchange contexts. The PS transfer rate found to be maximal at k_ON-PS_ = 3.7 µM^−1.^s^−1^ when PI(4)P was absent, slightly increased in the presence of PI(4)P (acceleration factor = 0.13). If k_ON-PS_ > 3.7 µM^−1^.s^−1^ the PS transfer rates were lower in a non-exchange situation but considerably higher if PI(4)P was present as counterligand. In contrast, for k_ON-PS_ < 3.7 µM^−1^.s^−1^, the PS transfer rate decreased toward zero, even if PI(4)P was present, and acceleration factors were almost null. In parallel, the PI(4)P transfer rates were found to be systematically higher in the presence of PS yet to a degree that depended on k_ON-PS_ values (**Figure 5b**, left panel). Finally, we noted that an ORD efficiently exchanges PS and PI(4)P if it has a higher affinity for PS than PI(4)P (k_ON-PS_ > k_ON-PI(4)P_, **Figure 5b**, left panel, grey area). These simulations again corroborated our experimental data. In non-exchange situations, saturated PS species, which are low-affinity ligands compared to 16:0/16:0-PI(4)P, are transferred at the fastest rates; yet these rates barely or marginally increase once PI(4)P is present. In contrast, unsaturated PS species, which are globally better ligands than 16:0/16:0-PI(4)P, are slowly transferred in non-exchange situations, but much faster when PI(4)P is present. In all cases, the PI(4)P transfer rate is unchanged or higher in the presence of PS.

To consolidate this analysis, we plotted the acceleration factors for PS and PI(4)P, simulated for different k_ON-PS_ and k_ON-PI(4)P_ values **(Figure 5–Figure Supplement 1a**, middle panels), against each other and compared the curves obtained with experimental factors shown in **Figure 1e**. By setting k_ON-PI(4)P_ at 40 or 75 µM^−1^.s^−1^, we obtained curves that follow the distribution of acceleration factors obtained with Osh6p and ORD8, respectively **(Figure 5c)**.

Finally, simulations performed with variable k_ON-PI(4)P_ values (10, 40, 75 or 100 µM^−1^.s^−1^) indicated that, when the ORD had a higher affinity for PI(4)P, the PI(4)P transfer rate decreased in a non-exchange context (dashed lines, **Figure 5–Figure Supplement 1a**) but increased to a greater extent in a situation of PS/PI(4)P exchange. Interestingly, a different picture emerged if the affinity of the ORD for PI(4)P was increased by ten, by lowering the k_OFF-PI(4)P_ value from 1 to 0.1 s^−1^ instead of increasing the k_ON-PI4P_ value from 10 to 100 µM^−1^.s^−1^ (**Figure 5–Figure Supplement 1b**): PI(4)P transfer rates were low in non-exchange conditions and only slightly higher in exchange conditions, for all tested k_ON-PS_ values. PS transfer was more rapid in exchange conditions with high k_ON-PS_ values. This resembled our data showing that unsaturated PI(4)P species were poorly transferred and exchanged for PS while PS transfer was enhanced by these PI(4)P species (**Figure 2**). Overall our model showed how variations in the capacity of the ORD to extract and deliver PS and PI(4)P could modify how it transfers and exchanges these lipids.

### Osh6p and ORD8 cannot use PI(4,5)P_2_ if PI(4)P is present in membranes

ORP5/8 have been suggested, notably based on *in vitro* data, to use PI(4,5)P_2_ instead of PI(4)P as a counterligand to supply the PM with PS^17^ but this conclusion is disputed^19^. To address this issue, we measured how ORD8 transferred PI(4)P and PI(4,5)P_2_ (with 16:0/16:0 acyl chains) from L_B_ liposomes that contained only one kind of PIP or, like the PM, both PIPs, to L_A_ liposomes. NBD-PH_FAPP_, which can detect PI(4,5)P_2_ in addition to PI(4)P^17^, was used as sensor. ORD8 transported PI(4,5)P_2_ more swiftly than PI(4)P (**Figure 6a,b**), as previously shown, but surprisingly, when both PIPs were in L_B_ liposomes, the transfer kinetics was comparable to that measured with PI(4)P alone.

**Figure 6.**
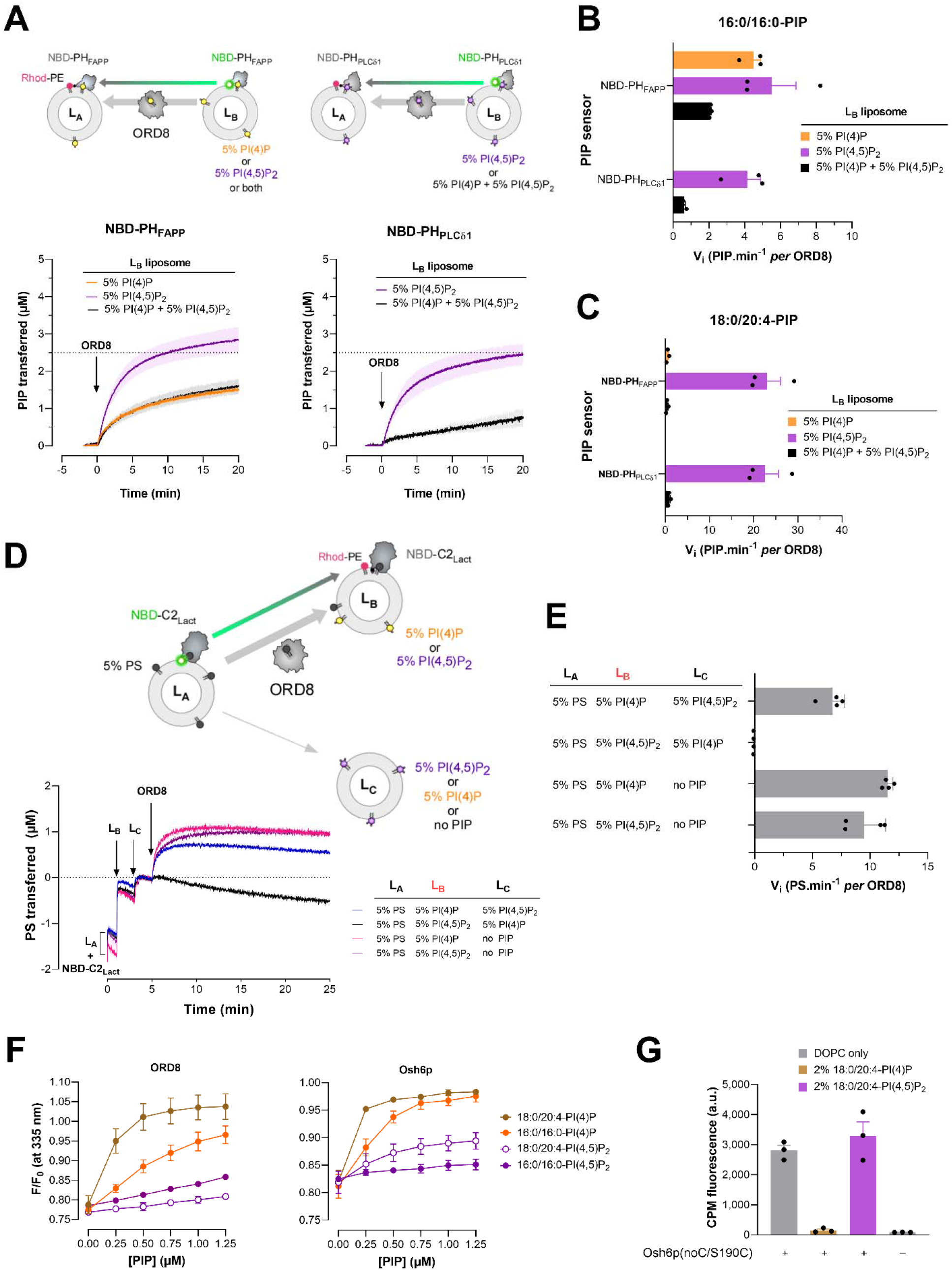
PIP selectivity of Osh6p and ORD8. **(A)** Intermembrane PI(4)P and PI(4,5)P_2_ transfer activity of ORD8 measured with donor L_B_ liposomes containing either PI(4)P or PI(4,5)P_2_ or both PIPs. L_B_ liposomes (200 µM total lipid concentration) made of DOPC and containing either 5% 16:0/16:0-PI(4)P or 5% 16:0/16:0-PI(4,5)P_2_ or both PIPs were mixed with NBD-PH_FAPP_ (250 nM) in HK buffer at 37°C. After 1 min, L_A_ liposomes, made of DOPC and containing 2% Rhod-PE (200 µM) were added to the reaction mix. After 3 minutes, ORD8 was injected (240 nM). Similar experiments were conducted with NBD-PH_PLCδ1_ (250 nM) and L_B_ liposomes containing only 5% 16:0/16:0-PI(4)P or also 5% 16:0/16:0-PI(4,5)P_2_. PIP transport was followed by measurement of the quenching of the fluorescence signal of NBD-PH_PLCδ1_ or NBD-PH_FAPP_ caused by the translocation of the lipid sensor from L_B_ to L_A_ liposomes and FRET with Rhod-PE. The signal was normalized in term of PI(4)P or PI(4,5)P_2_ amount delivered into L_A_ liposomes. The injection of ORD8 set time to zero. Each trace is the mean ± s.e.m. of independent kinetics (n = 3). **(B)** Initial 16:0/16:0-PIP transfer rates determined with NBD-PH_FAPP_ and NBD-PH_PLCδ1_. Data are represented as mean ± s.e.m. (error bars, n = 3). **(C)** Initial rates of 18:0/20:4-PIP transfer between membranes determined for ORD8 using NBD-PH_FAPP_ and NBD-PH_PLCδ1_ sensors. Data are represented as mean ± s.e.m. (error bars, n = 3). **(D)** L_A_ liposomes (200 µM total lipids), made of DOPC and 16:0/18:1-PS (95:5), were mixed with NBD-C2_Lact_ in HK buffer at 37°C. After 1 min, L_B_ liposomes (200 µM), consisting of 93% DOPC, 2% Rhod-PE and 5% 18:0/20:4-PI(4)P or 18:0/20:4-PI(4,5)P_2_, were injected. Two minutes later, a third population of liposomes (L_C,_ 200 µM) made of DOPC, doped or not with 5% 18:0/20:4-PI(4)P or 18:0/20:4-PI(4,5)P_2,_ was injected. ORD8 (240 nM final concentration) was injected two minutes later. PS delivery into L_B_ liposomes was followed by measurement of the quenching of the fluorescence of NBD-C2_Lact_ provoked by its translocation from L_A_ to L_B_ liposomes and FRET with Rhod-PE. The signal was normalized in term of PS amount transferred into L_B_ liposomes. **(E)** Initial rates of ORD8-mediated PS transfer into acceptor L_B_ liposomes, as a function of the presence of PI(4)P and PI(4,5)P_2_ in acceptor L_B_ and/or L_C_ liposomes. Data are represented as mean ± s.e.m. (error bars, n = 4). **(F)** Competition assay. DOPC/NBD-PS liposomes (98:2, 100 µM total lipid) were added to ORD8 or Osh6p (200 nM) in HK buffer at 30°C. The sample was excited at 280 nm and the emission was measured at 335 nm. Incremental amounts of liposome, containing 5% PI(4)P or PI(4,5)P_2_ were injected to the reaction mix. The signal was normalized considering the initial F_max_ fluorescence, prior to the addition of NBD-PS-containing liposomes, and the dilution effect due to liposome addition. Data are represented as mean ± s.e.m. (n = 3). **(G)** Accessibility assay. CPM (4 µM) was mixed with 400 nM Osh6p(noC/S190C) in the presence of pure DOPC liposomes or liposomes doped with 2% 18:0/20:4-PI(4)P or 18:0/20:4-PI(4,5)P_2_. Intensity bars correspond to the fluorescence measured 30 min after adding CPM (n = 3).

To understand why, we specifically measured the transfer of PI(4,5)P_2_ with a sensor based on the PH domain of the phospholipase C-δ1 (PH_PLCδ1_), which has a high affinity and specificity for the PI(4,5)P_2_ headgroup^42^. This domain was reengineered to include, near its PI(4,5)P_2_-binding site^43^, a unique solvent-exposed cysteine (C61) to which a NBD group was attached (**Figure 6 – Figure Supplement 1a**). In flotation assays this NBD-PH_PLCδ1_ construct associated with liposomes doped with PI(4,5)P_2_ but not with liposomes only made of DOPC or containing either PI or PI(4)P (**Figure 6 – Figure Supplement 1b**). Fluorescence assays also indicated that NBD-PH_PLCδ1_ bound to PI(4,5)P_2_-containing liposomes, as its NBD signal underwent a blue-shift and a 2.2-fold increase in intensity (**Figure 6 – Figure Supplement 1c**). A binding curve was established by measuring this change as a function of the incremental addition of these liposomes (**Figure 6 – Figure Supplement 1c**). In contrast, no signal change occurred with PI(4)P-containing liposomes, indicative of an absence of binding (**Figure 6 – Figure Supplement 1c,d**). NBD-PH_PLCδ1_ was thus suitable to detect PI(4,5)P_2_ but not PI(4)P. It was substituted for NBD-PH_FAPP_ to measure to what extent ORD8 specifically transported PI(4,5)P_2_ from L_B_ liposomes, containing or not PI(4)P, to L_A_ liposomes. Remarkably, we found that ORD8 efficiently transferred PI(4,5)P_2_ but only if PI(4)P was absent (**Figure 6a,b**). Similar conclusions were reached using each PIP sensor by assaying ORD8 with PI(4)P and PI(4,5)P_2_ ligands with 18:0/20:4 acyl chains (**Figure 6c, Figure 6 – Figure Supplement 2a**), and Osh6p using PIPs with a 16:0/16:0 composition (**Figure 6 – Figure Supplement 2b, c**). These data indicate that ORP8 and Osh6p preferentially extract PI(4)P from a membrane that contains both PI(4)P and PI(4,5)P_2_, suggesting that they use PI(4)P rather than PI(4,5)P_2_ in exchange cycles with PS at the PM.

To address this possibility *in vitro*, we devised an assay with three liposome populations (**Figure 6d, e)** to examine whether ORP8 delivers PS in a PI(4)P-containing membrane or in a PI(4,5)P2-containing membrane. First, L_A_ liposomes doped with 5% PS were mixed with NBD-C2_Lact_. Then L_B_ liposomes, containing 5% PI(4)P and 2% Rhod-PE, and L_C_ liposomes, only made of PC, were successively added. Injecting ORD8 provoked a quenching of the NBD signal, indicating that the C2_Lact_ domain moved onto the L_B_ liposomes. The signal normalization indicated that ∼1 µM of PS was transferred to L_B_ liposomes. Equivalent data were obtained with L_C_ liposomes doped with PI(4,5)P_2_, suggesting that this lipid has no influence on the PI(4)P-driven transfer of PS to L_B_ liposomes mediated by ORD8. We performed mirror experiments with L_B_ liposomes that contained PI(4,5)P_2_ and L_C_ liposomes with or without PI(4)P. Remarkably, PS was transferred to L_B_ liposomes but not if L_C_ liposomes contained PI(4)P. We concluded that ORP8 selectively delivers PS in a compartment that harbors PI(4)P if PI(4,5)P_2_ is present in a second compartment. This suggests that PI(4)P, and not PI(4,5)P_2_, is used by ORP5/8 to transfer PS intracellularly.

These observations suggest that ORD8 and Osh6p have a lower affinity for PI(4,5)P_2_ than for PI(4)P. Confirming this, the NBD-PS-based competition assay showed that each protein barely bound to PI(4,5)P_2_ compared to PI(4)P (with 16:0/16:0 or 18:0/20:4 acyl chains **Figure 6f)**. Likewise, TSAs indicated that Osh6p incubated with liposomes containing 16:0/16:0- and 18:0/20:4-PI(4)P or PI(4,5)P_2_ was loaded with and stabilized by PI(4)P but not PI(4,5)P_2_ (**Figure 6 – Figure Supplement 2d,e**). Finally, we evaluated the conformational state of Osh6p in the presence of each PIP. To this end, we used a version of the protein, Osh6p(noC/S190C), which has a unique cysteine at position 190; this residue is solvent-exposed only if the molecular lid that controls the entrance of the binding pocket of the protein is open^44^. This construct was added to liposomes devoid of PIPs or containing 2% PI(4)P or PI(4,5)P_2_. Then, 7-diethylamino-3-(4’-maleimidylphenyl)-4-methylcoumarin (CPM), a molecule that becomes fluorescent only when forming a covalent bond with accessible thiol, was added to each sample. After a 30-min incubation, a high fluorescence signal was measured with Osh6p(noC/S190C) mixed with PC liposomes, indicating that the protein remained essentially open over time (**Figure 6g**). In contrast, almost no fluorescence was recorded when the protein was incubated with PI(4)P-containing liposomes, indicating that it remained mostly closed, as previously shown^44^. Remarkably, a high signal was obtained with liposomes doped with 2% PI(4,5)P_2_, indicating that Osh6p remained open as observed with pure PC liposomes. Altogether, these data suggest that Osh6p/7p and ORP5/8 have a low affinity for PI(4,5)P_2_ compared to PI(4)P, likely as they cannot form stable and closed complexes with this lipid.

### Sterol abundance in membrane does not enhance PS delivery and retention

Like PS, sterol is synthesized in the ER and enriched in the PM, where it constitutes 30-40% of all lipids^45^. PS is thought to associate laterally with sterol, thus retaining sterol in the inner leaflet of the PM and controlling its transbilayer distribution^30,31^. However, it was not known whether sterol stabilizes PS and thereby aids ORP/Osh proteins to accumulate PS in the PM. To explore this possibility, we measured *in vitro* the speed at which Osh6p transported PS from L_A_ liposomes to L_B_ liposomes, containing or not 30% cholesterol or ergosterol, and doped or not with 5% PI(4)P. These assays were performed using 16:0/18:1-PS and 16:0/18:1-PI(4)P, as in yeast, these predominant PS and PI(4)P species are thought to preferentially populate sterol-rich nanodomains in the PM^22^. However, we observed that the transfer of PS was not markedly impacted by higher contents of sterol in L_B_ liposomes in non-exchange and exchange conditions (**Figure 7a, Figure 7 – Figure Supplement 1a**). We then replaced 16:0/18:1-PS in our assays with 18:0/18:1-PS shown to segregate with cholesterol *in vitro*^31^. Yet, with L_B_ liposomes containing 0, 30 or even 50% cholesterol, no change was seen in the 18:0/18:1-PS transfer rate in a non-exchange context. If PI(4)P was present in L_B_ liposomes, PS was transferred more rapidly but at a slightly lesser extent when L_B_ liposomes also contained 50% cholesterol (**Figure 7b,c**). This suggests that sterols do not favor PS transfer, possibly as they are dispensable for PS retention. To examine more extensively how cholesterol controls the retention of PS in the complex context of the PM, we used a GFP-C2_Lact_ probe to examine whether the steady-state accumulation of PS in the PM of HeLa cells was impacted when cholesterol was depleted from the PM. This depletion was achieved by treating the cells for 24 h with U18666A, a compound that blocks lysosomal-to-PM sterol movement^31,46,47^. Such a treatment lowers sterol levels without provoking the remodeling of the PM that occurs following a faster and acute sterol removal^30^. Using the sterol-sensor mCherry-D4, we detected sterol in the PM of untreated cells but not in U18666A-treated cells **(Figure 7d,e)**. The same results were obtained with a mCherry-D4H construct (carrying the D434S mutation) even if it can detect lower sterol density^31^, confirming that the PM was highly deprived in sterol (**Figure 7 – Figure Supplement 1b**). However, no change was seen in the distribution of PS, which remained in the PM. This was ascertained by measuring the relative distribution of GFP-C2_Lact_ between the PM and the cytosol, in treated and untreated cells, using Lyn11-FRB-mCherry as a stable PM marker and internal reference (**Figure 7d, Figure 7 – Figure Supplement 1c**). We conclude that the presence of sterol in the PM is not essential for PS retention and thus for PS/PI(4)P exchange.

**Figure 7.**
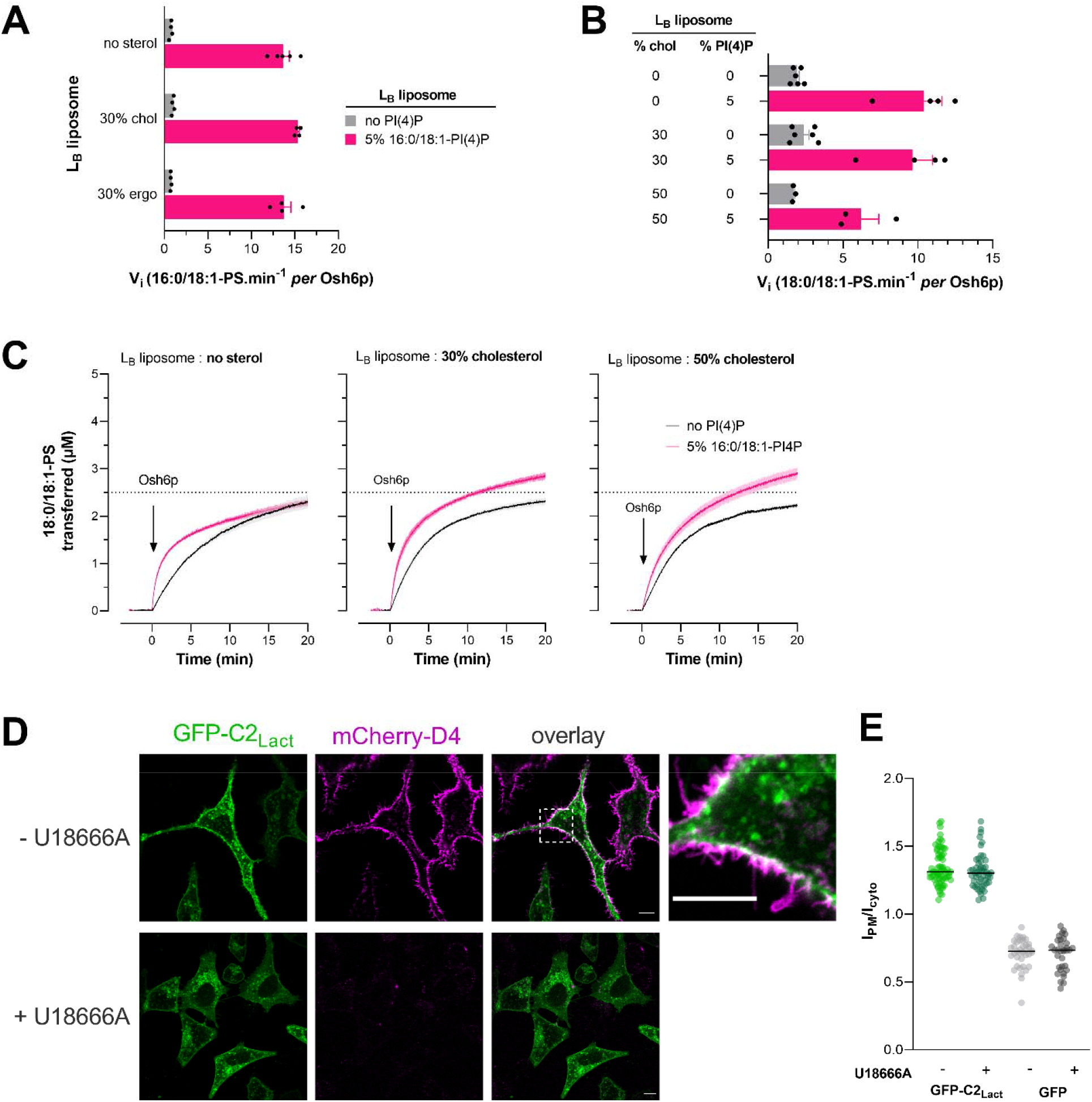
Influence of the sterol content of acceptor membranes on PS delivery. **(A)** Initial rates of 16:0/18:1-PS transfer measured with Osh6p (200 nM) at 30°C from L_A_ (5% PS) to L_B_ liposomes with different bulk lipid compositions and containing or not 5% 16:0/18:1-PI(4)P (at the expense of PC). Data are represented as mean ± s.e.m. (non-exchange condition, n = 4-5, exchange condition, n = 4). **(B)** Initial rates of 18:0/18:1-PS transfer from L_A_ to L_B_ liposomes without or with 30% or 50% cholesterol, and containing or not 5% 16:0/18:1-PI(4)P. Data are represented as mean ± s.e.m. (n = 3-6). **(C)** Experiments are similar to those shown in **(A)** except that 16:0/18:1-PS was replaced by 18:0/18:1-PS in L_A_ liposomes. L_B_ liposomes contained 0, 30 or 50% cholesterol. Each trace is the mean ± s.e.m. of kinetics recorded in independent experiments (n = 3-6). **(D)** Confocal images of live GFP□C2_Lact_□expressing HeLa cells (green) treated or not with U18666A for 24 h at 37°C. Cholesterol in the PM was detected by incubating the cells for 10 min with mCherry-D4 (purple) at room temperature and then washed with medium prior to imaging. The overlay panel shows merged green and magenta images. The top-right image corresponds to a higher magnification (5.2 ×) image of the area outlined in white. Scale bars: 10 µm. **(e)** Quantification of the ratio of GFP□C2_Lact_ signals at the PM to the cytosolic GFP□C2_Lact_ signals, as assessed by wide-field microscopy and line scan analysis. GFP□C2_Lact_□expressing HeLa cells with or without U18666A treatment (2.5 µg/mL for 24 h at 37°C) as shown in **(D)** (mean ± s.e.m., n = 68 cells for non-treated cells and n = 51 cells for treated cells; data are pooled from four independent experiments for each condition). Control experiments were done with GFP□expressing HeLa cells (mean ± s.e.m., n = 37 cells for non-treated cells and n = 33 cells for treated cells; data are pooled from two independent experiments for each condition).

## Discussion

LTPs have been discovered and studied for more than 40 years, yet few studies have explored how their activity depends on the nature of the lipid acyl chains. A few kinetic studies have shown that a nonspecific-LTP transfers shorter PC more easily than long PC^48^, that a glycolipid transfer protein (GLTP) preferentially transports short glucosylceramides^49^ and that the ceramide transfer protein (CERT) is active with ceramide species whose length does not exceed the size of its binding pocket^50,51^. Moreover, the link between the activity of LTPs and their affinity for lipid ligands remained largely obscure. Here, we measured how fast LTPs transfer saturated *vs* unsaturated lipids between membranes, both in a situation of simple transfer and in a situation of exchange with a second ligand, and we measured their relative affinity for these lipids. Our investigations used approaches that detect the transfer and binding capacities of LTPs with unmodified lipid ligands (i.e., without bulky fluorophores), thus with an unprecedented level of accuracy. This study offers novel insights into PS/PI(4)P exchangers and, by identifying how the activity of LTPs relates to their affinity for lipid ligands, provides general rules that serve to better analyze lipid transfer processes.

Our kinetic measurement indicated that overall ORD8 transfers PS and PI(4)P more slowly than Osh6p does. This might be due to structural differences between the two proteins or to the fact that ORD8 only functions optimally in the context of the full-length ORP8 or between closely-apposed membranes. Apart from this, Osh6p and ORD8 responded similarly to the same changes in the PS and PI(4)P acyl chain structures. In particular, they transferred these lipids quite differently depending on whether they were saturated or not. This seems to be mainly because these LTPs have a lower affinity for saturated than unsaturated lipids. Remarkably the presence of only double bond in one of the acyl chains of the ligand is sufficient to significantly change their behavior. Our kinetic model suggests that the affinity of PS/PI(4)P exchangers for lipid ligands and its capacity to transfer them is governed by the extraction step (reflected by the k_on_ values). An early study of large series of PE and PC species showed that increasing the degree of unsaturation in the acyl chains of phospholipids increases the rate at which they spontaneously desorb from the membrane^52^. One might therefore posit that the intrinsic propensity of PS species to move out of the membrane determines how they are captured and transferred by ORP/Osh proteins. However, several data suggest that the intrinsic tendency of PS species to leave the membrane cannot explain, or only very partially, why these lipids are more or less easily captured by Osh6p and ORD8.

First, considering the spontaneous desorption rates measured with other phospholipids^52^, we should have obtained the exact same results with 18:0/18:1-PS as with 18:1/18:0-PS, which was not the case, notably in binding assays. Secondly, 16:0/16:0-PS should have been as good a ligand as an unsaturated species such as 16:0/18:1-PS and 18:1/18:1-PS whereas shorter saturated PS species (12:0/12:0 and 14:0/14:0) should have been better. Also, similar results should have been obtained with 16:0/16:0-PI(4)P and 16:0/18:1-PI(4)P. Structural analyses revealed that the sn-1 acyl chain of PS or PI(4)P is deeply inserted in the binding pocket, and that their sn-2 acyl chain is twisted, pointing its end to the lid that closes the pocket^12,13^. These structural constraints along with other parameters, such as the intrinsic dynamics of lipid species in a bilayer, might govern how they are captured and stabilized by ORP/Osh proteins. In the future, studies addressing these hypotheses at the atomic level will be of great interest to understand what imparts these LTPs with such an enigmatic selectivity for unsaturated lipids. For instance, solving the structure of Osh6p or ORD8 in complex with various lipid species along with molecular dynamics (MD) simulations^53–55^ might shed light on how lipid species are stabilized inside the binding pocket, released into or extracted from the membrane.

Kinetic data obtained with unsaturated lipids that correspond to or resemble PS and PI(4)P species that are prominent in yeast and human cells, provided hints on the activity of these LTPs in cellular context, which remains difficult to obtain *in situ*. Osh6p slowly transfers 16:0/18:1-PS and 16:0/18:1-PI(4)P, the most abundant yeast PS and PI(4)P species^22,24,28,36^, under non-exchange conditions and ten times faster under exchange condition, and with the same initial velocity (∼14 lipid.min^−1^). This suggests that, at the ER/PM interface, the transfer of PS by Osh6p/7p in yeast is highly dependent on the synthesis of PI(4)P in the PM and is tightly coupled with the transfer of PI(4)P in the opposite direction. Another PS species, 18:0/18:1-PS paired with 16:0/16:0-PI(4)P or 16:0/18:1-PI(4)P efficiently completes fast exchange, as does 18:1/18:1-PS with 16:0/16:0-PI(4)P. These PS species do not exist in yeast but resemble 16:0/18:1-PS and the few other major yeast PS species (16:0/16:1, 16:1/18:1, 16:1/16:1), which contain at least one monounsaturated acyl chain. In comparison, PS species with one or two di-unsaturated chains seemed less suitable for exchange processes. An equivalent analysis with PI(4)P remains difficult to conduct as much fewer PI(4)P species were assayed. The fact that polyunsaturated PI(4)P, the major brain PI(4)P species, exchanges poorly with 16:0/18:1-PS does not mean that Osh6p is specifically designed for yeast monounsaturated PI(4)P. Indeed, slow transfers were measured when assaying polyunsaturated PI(4)P with ORD8. Collectively our data suggest that Osh6p can efficiently exchange unsaturated PS and PI(4)P species at the ER/PM interface, supporting the notion that Osh6p/7p regulate a pool of unsaturated PS and PI(4)P that together with ergosterol form functional lipid nanodomains^22^. However, our investigation suggests that the delivery of PS in the PM, and its exchange for PI(4)P is not specifically promoted by the presence of ergosterol. Finally, our data suggest that the massive increase in the proportion of 16:0/18:1-PS observed between the ER membrane and the PM^28^ (from 30 to 60% compared to the other PS species) is not caused by a preferential transfer of this species by Osh6p/7p.

Analyzing the speed at which ORD8 transfers 16:0/18:1 and 18:0/18:1-PS, the prevalent PS species in human cells, we draw conclusions similar to those obtained with Osh6p. These PS species are slowly transferred under non-exchange conditions and faster under exchange conditions to an extent that depends on the PI(4)P species used as counterligand (16:0/16:0, 18:1/18:1 or 18:0/20:4-PI(4)P). This suggests that the PS transfer activity of ORP5/8 intimately depends on PI(4)P but is likely not influenced by its unsaturation degree, a parameter that varies between cells in tissues and cultured transformed cells. When exchange is possible, the transfer of 18:0/20:4-PI(4)P and 18:1/18:1-PI(4)P is faster but remains, intriguingly, much slower than the PS transfer, suggesting a weak coupling between the two transfer processes. This presumably arises from the high affinity of the ORD for these PI(4)P species, as suggested by our kinetic model. Possibly, the hydrolysis of PI(4)P by Sac1 at the ER enhances through mass action the transfer of PI(4)P from the PM to the ER by ORP5/8 while facilitating the extraction of PS from this organelle, and thus improves the coupling between PS and PI(4)P transfer at the ER/PM interface. Of note, PI(4)P hydrolysis is mandatory for OSBP to execute the sterol/PI(4)P exchange as OSBP has a much higher affinity for PI(4)P than for sterol^56^. Last we found *in vitro* that cholesterol does not promote the delivery of 18:0/18:1-PS in an acceptor membrane and the retention of PS in the PM of human cells.

It was unclear whether ORP5/8 could use PI(4,5)P_2_ instead of PI(4)P as a counterligand^14,17,19^. Corroborating previous observations^17^, we measured that Osh6p and ORD8 transferred PI(4,5)P_2_ between liposomes more rapidly than PI(4)P. However, when PI(4)P and PI(4,5)P_2_ both resided in the same donor membrane, only PI(4)P was transferred to acceptor membranes, for all tested acyl chain compositions. Moreover, only PI(4)P was used as counterligand for the transfer of PS when both PIPs were present. In fact, Osh6p and ORD8 show a much lower affinity for PI(4,5)P_2_ than for PI(4)P. This likely relates to the fact that PI(4,5)P_2_, contrary to PI(4)P (or PS), cannot be entirely buried in the pocket and capped by the lid. This hypothesis was suggested by structural analyses of ORP1 and ORP2 in complex with PI(4,5)P_2_^34,35^ and confirmed by our *in vitro* assays. Because PI(4)P and PI(4,5)P_2_ co-exist in the PM, this strongly suggests that only PI(4)P is used by ORP5 or ORP8 for the exchange with PS in the cell. Our data also suggest that ORP1, ORP2 and ORP4L^57^ cannot trap PIPs other than PI(4)P in cells or only if they operate on organelle membranes devoid of PI(4)P.

Quite interestingly, the comparison between the transfer rates and affinity determined with Osh6p and ORD8 for various lipid species allows us to infer general rules that can serve to better understand the cellular activity of LTPs. A first lesson is that, in a simple transfer process between membranes, a low-affinity ligand can be transferred more rapidly down its concentration gradient than a high-affinity ligand. This was observed when comparing saturated PS with unsaturated PS or PI(4,5)P_2_ with PI(4)P. Presumably, as suggested by our kinetic model, it lies in the fact that a low affinity recognition process can be detrimental when the ligand is extracted from a donor membrane but an advantage in the transfer process, notably by preventing any re-extraction of the ligand from the acceptor membrane.

The picture is different with a membrane system of higher complexity that reconstitutes a cellular context more faithfully. In exchange conditions, the ability of Osh6p and ORD8 to saturated PS was poorly enhanced or often inhibited when PI(4)P was present as a counterligand, because these LTPs have a higher affinity for the latter. When PI(4)P and PI(4,5)P_2_ were present in the same donor membrane, these LTPs preferentially transferred PI(4)P, for which they have the highest affinity, to acceptor membranes. In an even more complex system where PS, PI(4)P and PI(4,5)P_2_ were present each in distinct liposome populations, mimicking three cellular compartments, ORD8 used PI(4)P as a counterligand to transfer PS between two membranes.

These observations have important implications. They suggest first that some caution must be exercised when analyzing *in vitro* data using membranes of low compositional complexity: measuring a fast transfer rate for a given lipid ligand and an LTP does not necessarily mean that this ligand is the true cellular cargo of the LTP. When considering a cellular context, one can assume that a mere lipid transporter preferentially recognizes and transfers its high-affinity lipid ligand. This potentially implies, as suggested *in vitro*, a lower speed of transfer but at the benefit of a higher accuracy as no fortuitous ligand can be taken. However, this limitation in terms of speed is lifted if the LTP can exchange this high-affinity ligand for a second one, as measured with Osh6p and ORP8 using unsaturated PS and PI(4)P. This can primarily be explained by the fact that this second ligand prevents the re-extraction by the LTPs of the other ligand from its destination compartment, which improves its net delivery. These observations on Osh6p and ORP8 confirm very first data on the sterol/PI(4)P exchange capacity of Osh4p^33^. Of note, our experiment and models suggest that an optimal exchange occurs, *i.e.* with similar PS and PI(4)P transfer rates along opposite directions, when a lipid exchanger has a similar affinity for each ligand. Interestingly, the experiments with PS, PI(4)P and PI(4,5)P_2_ even suggest that this exchange process can channel the lipid flux between two membrane-bound compartments if there are more than two compartments, such as in a cell. Collectively, our study supports the notion that lipid exchange processes are mechanisms that ensure fast, accurate and directional transfer of lipids between organelles.

## Acknowledgments

We wish to thank Pr. Pietro de Camilli for providing the plasmid coding for the GST-ORD8 construct, Dr. Enrique Castano for the plasmid coding for the GST-PH_PLCδ1_ and Dr. Fabien Alpy for the plasmid coding for the mCherry-D4-His_6_. We thank Dr. Frédéric Brau for his help in image analysis. We are grateful to Ms. Y. Van Der Does for her careful corrections and proofreading of the manuscript. This work was supported by the CNRS and by a grant from the *Agence Nationale de la Recherche* (ANR-16-CE13-0006). NFL was supported by a fellowship from the *Ministère de l’Enseignement Supérieur, de la Recherche et de l’Innovation*.

## Author contributions

G.D. designed and supervised research. S.I., N-F. L., M.M., N.F. and G.D. carried out site-directed mutagenesis, produced, purified and labeled all the recombinant proteins of this study. S.I., N-F.L., N.F. V.D. and G.D. performed the *in vitro* experiments. M.M performed the cellular experiments. G.D., S.I., N-F.L, V.D. and M.M. analyzed the data. G.D wrote the manuscript. All of the authors discussed the results and commented on the manuscript.

## Supplementary Figures

**Figure 1-Figure Supplement 1.**
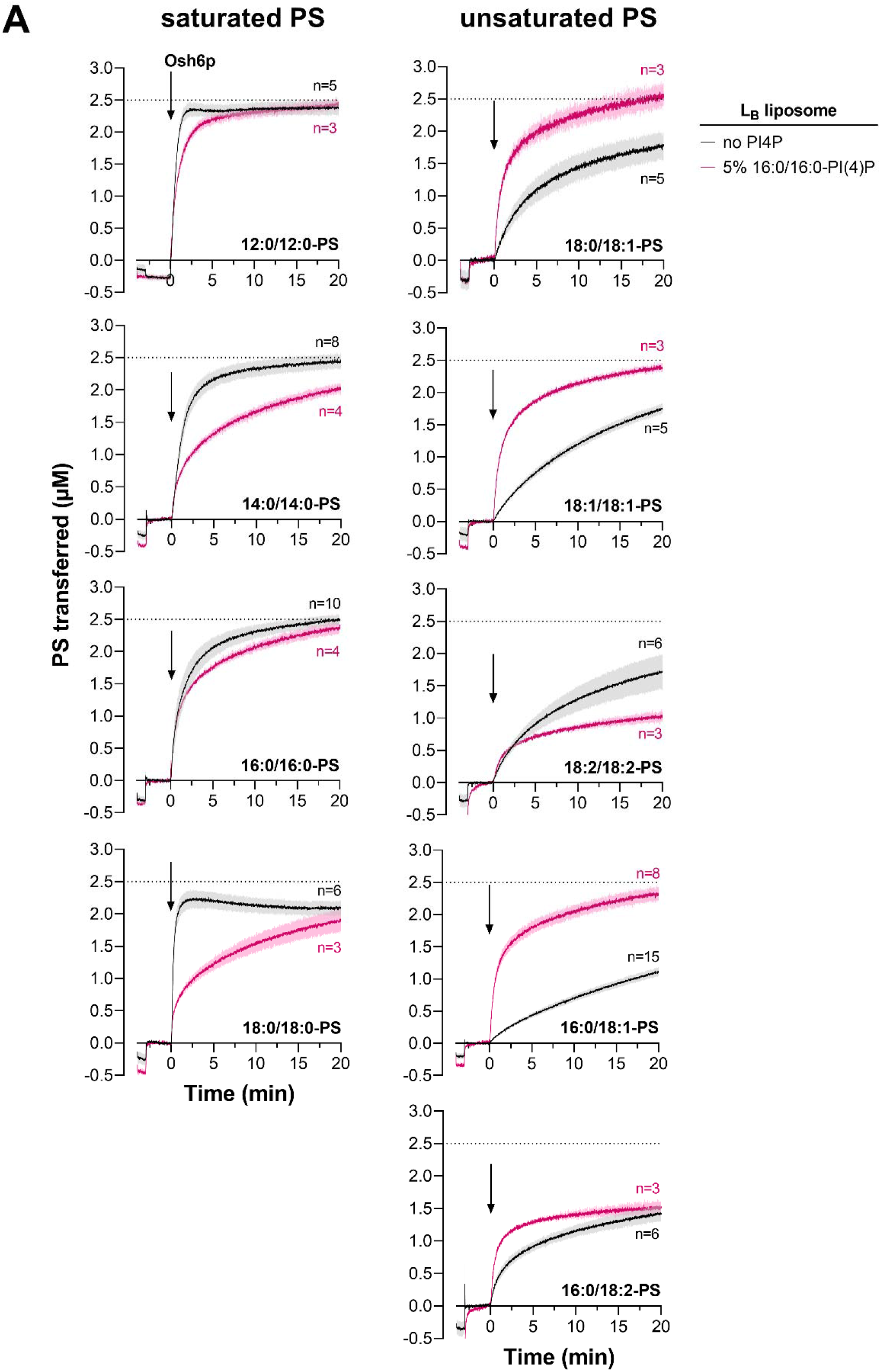
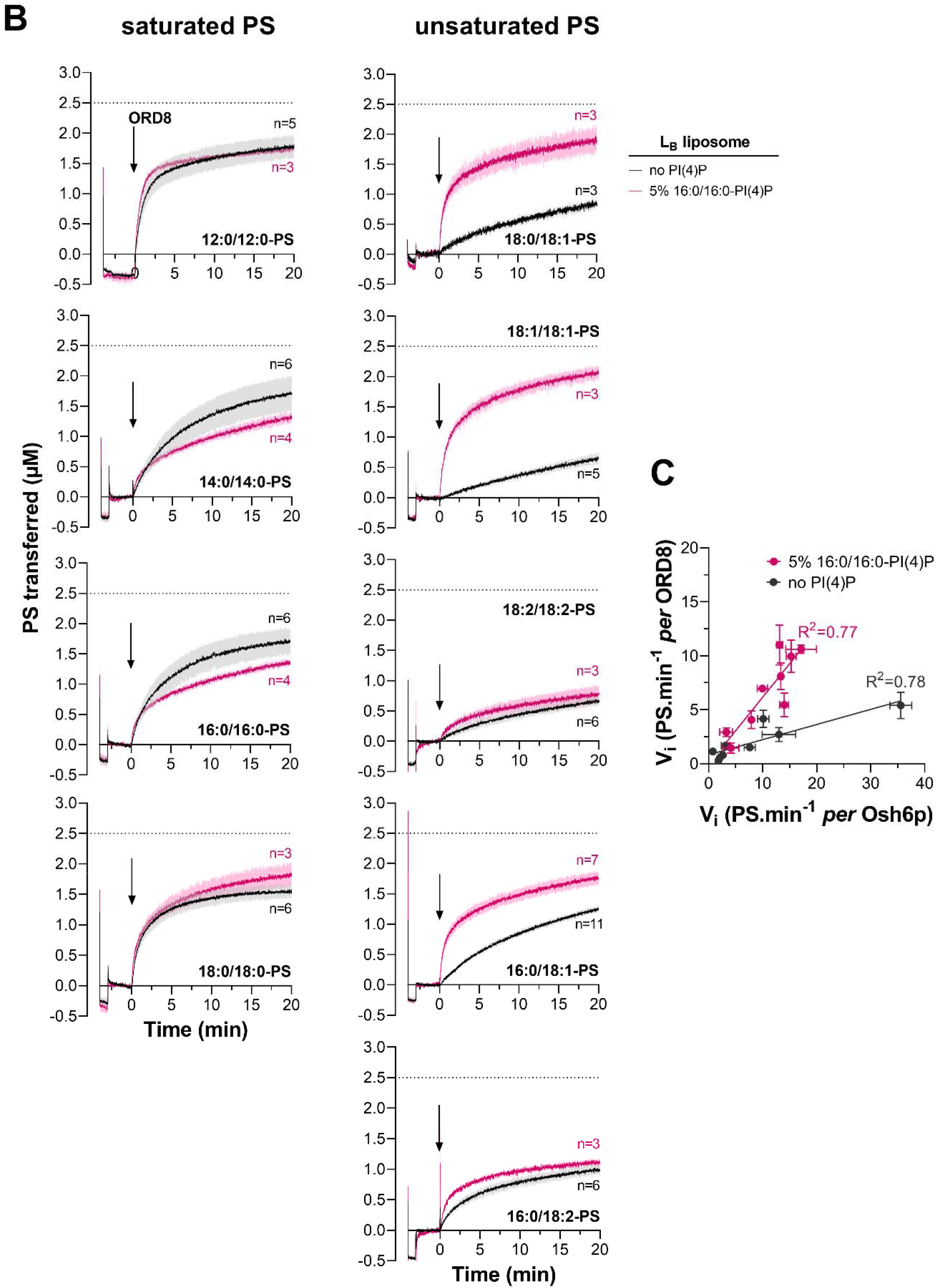
Kinetics of PS transfer measured with Osh6p and ORD8 with different PS species and L_B_ liposomes containing or not PI(4)P. (**A**) DOPC liposomes (200 µM total lipid, L_A_) containing 2% Rhod-PE and 5% PS were mixed with NBD-C2_Lact_ (250 nM) at 30°C. After one minute, DOPC liposomes (200 µM lipids, L_B_) containing or not 5% 16:0/16:0-PI(4)P were added. After 3 min, Osh6p (200 nM) was injected. Dark trace: without PI(4)P, pink trace: with PI(4)P. Each trace is the mean ± s.e.m. of kinetics recorded in independent experiments (n = 3-15). A dashed line indicates the level of PS in L_B_ liposomes that should be observed if the liposomes are fully equilibrated. (**B**) Similar experiments were carried out with ORD8 (240 nM) at 37°C. Each curve is the mean ± s.e.m. of curves recorded in independent experiments (n = 3-11). **(c)** Initial PS transfer rates measured with Osh6p *vs* those measured with ORD8 in the presence of L_B_ liposomes enriched or not with 5% 16:0/16:0-PI(4)P.

**Figure 1-Figure Supplement 2.**
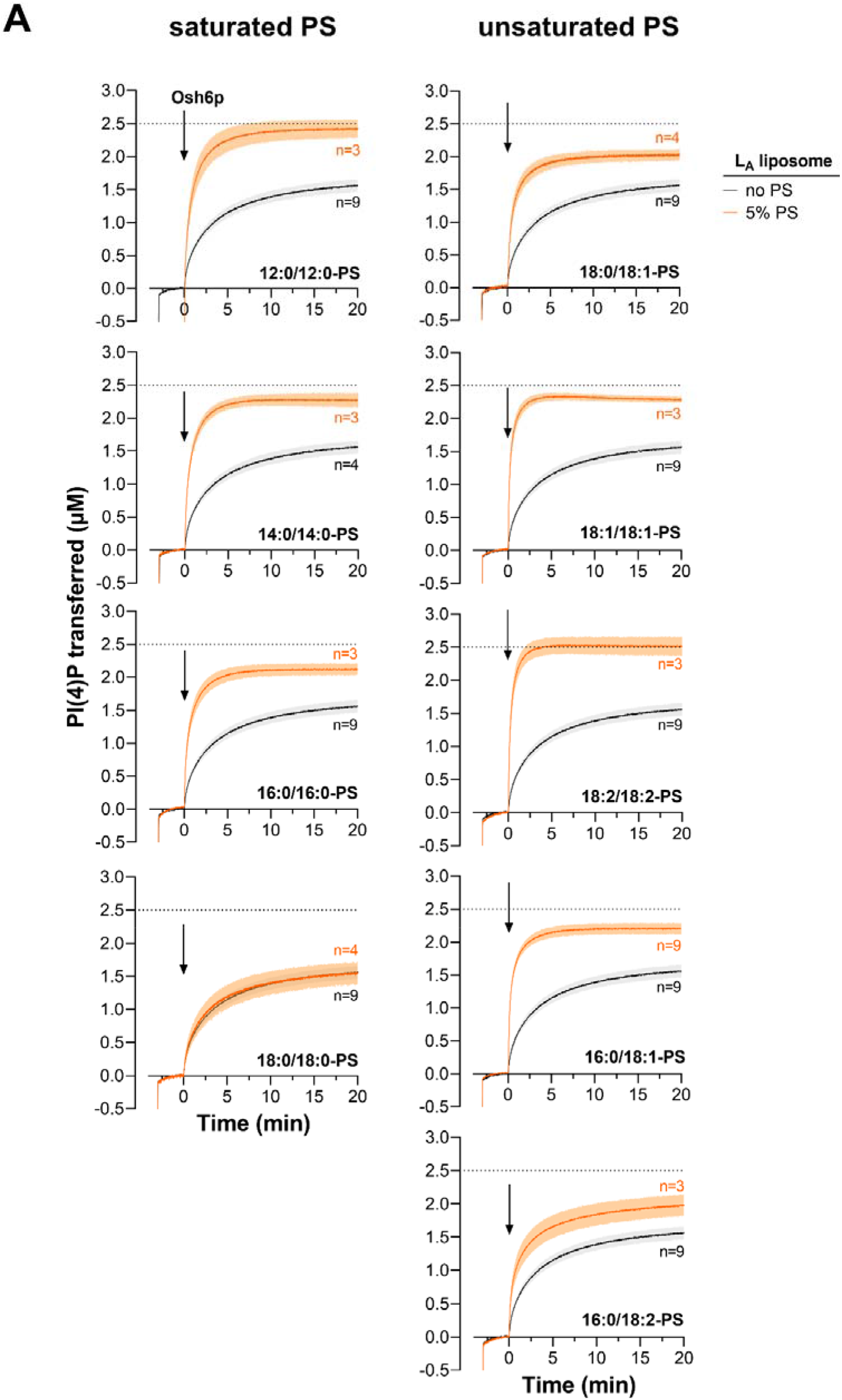
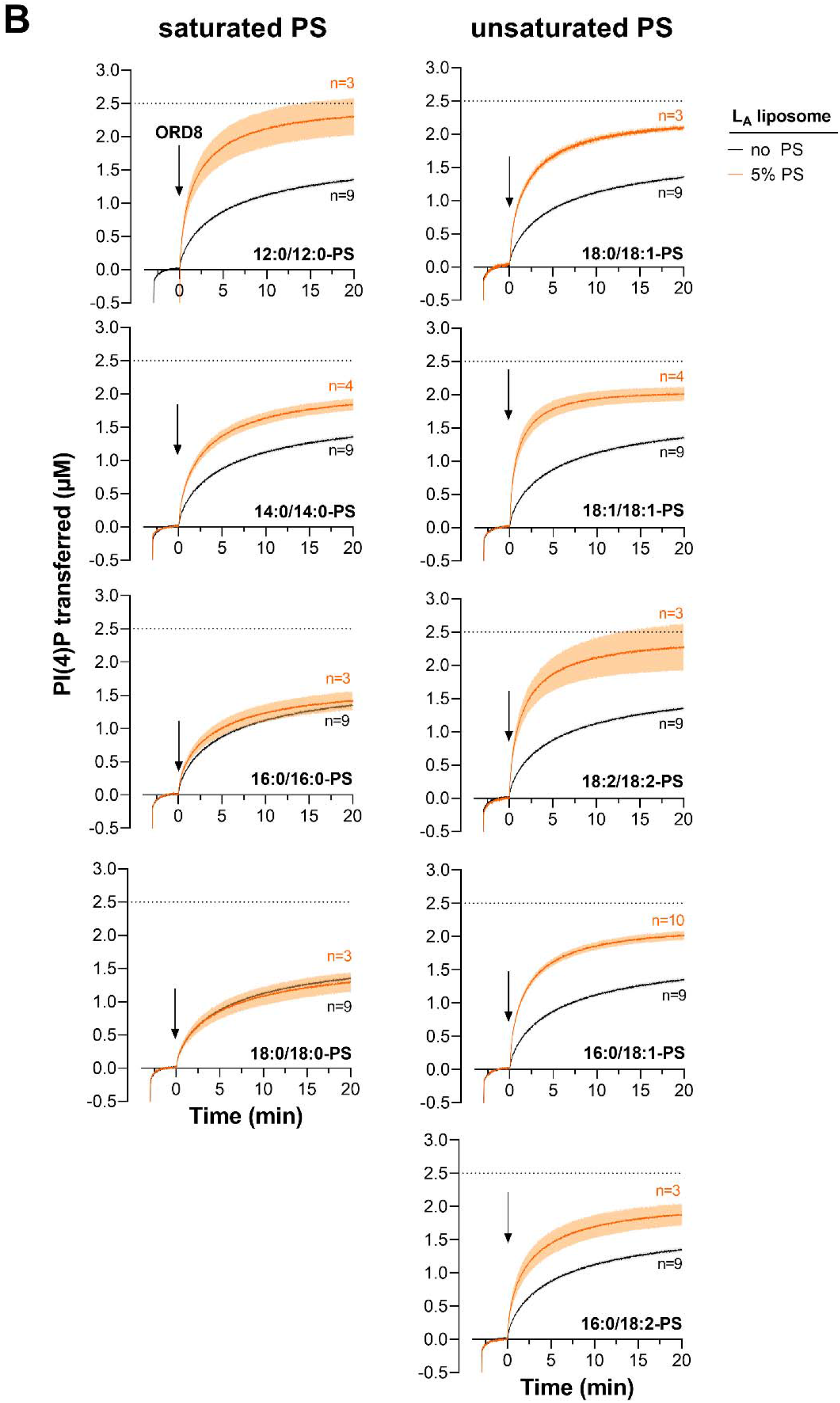
Kinetics of PI(4)P transfer by Osh6p and ORD8 in the context of PS/PI(4)P exchange depends on the length and unsaturation degree of PS species. **(A)** DOPC liposomes (200 µM lipids, L_B_) containing 5% 16:0/16:0-PI(4)P were mixed with NBD-PH_FAPP_ (250 nM) at 30°C. After one minute, DOPC liposomes (200 µM total lipid, L_A_) containing 2% Rhod-PE and 5% PS were added. After 3 min, Osh6p (200 nM) was injected. Dark trace: without PS, orange trace: with PS. Each trace is the mean ± s.e.m. of independent kinetics (n = 3-9). (**B**) Similar experiments were performed with ORD8 (240 nM) at 37°C. Each curve is the mean ± s.e.m. of independent kinetics traces (n = 3-10). A dashed line indicates the PI(4)P amount that should be transferred in L_B_ liposomes at equilibrium.

**Figure 2-Figure Supplement 1.**
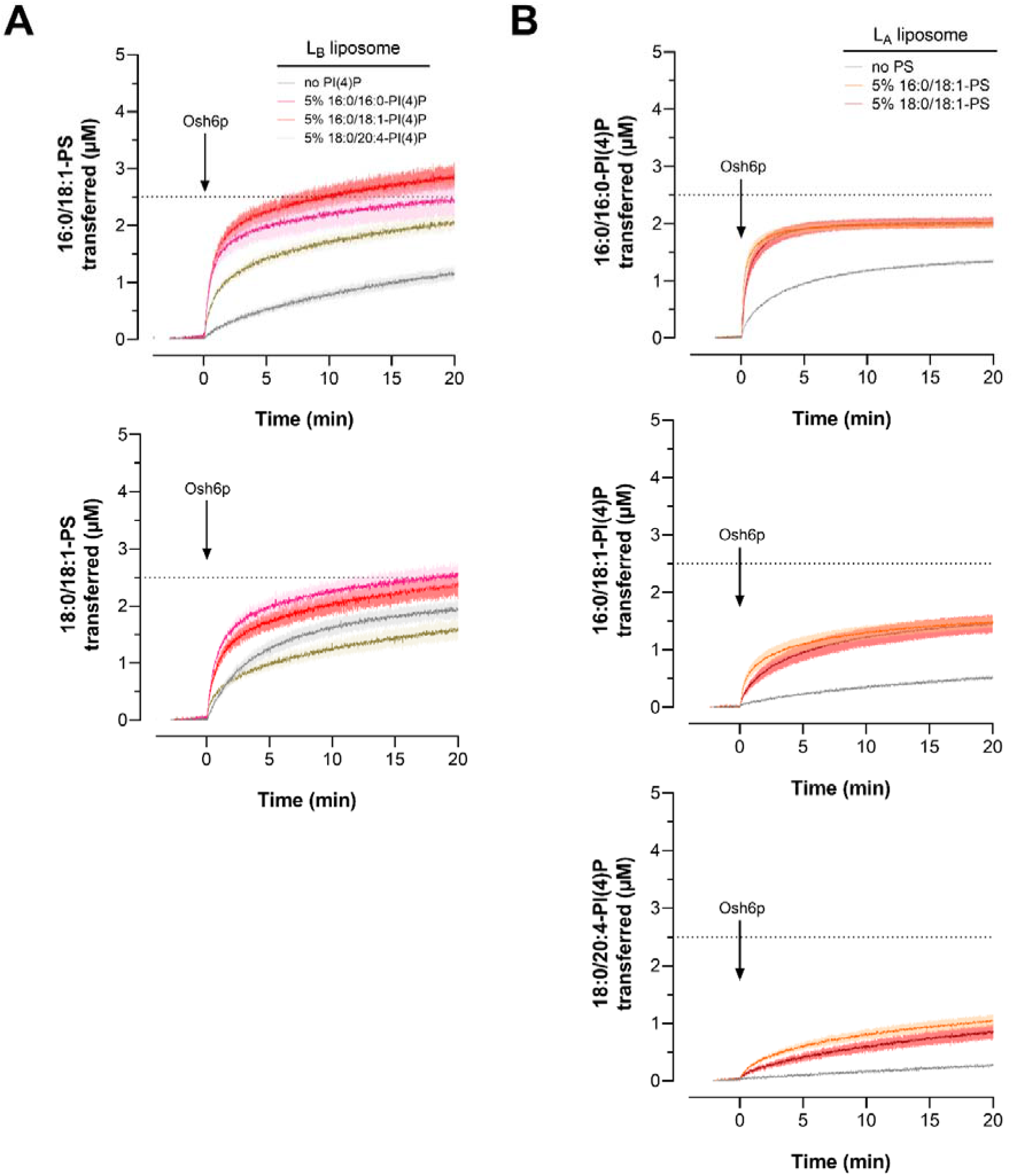
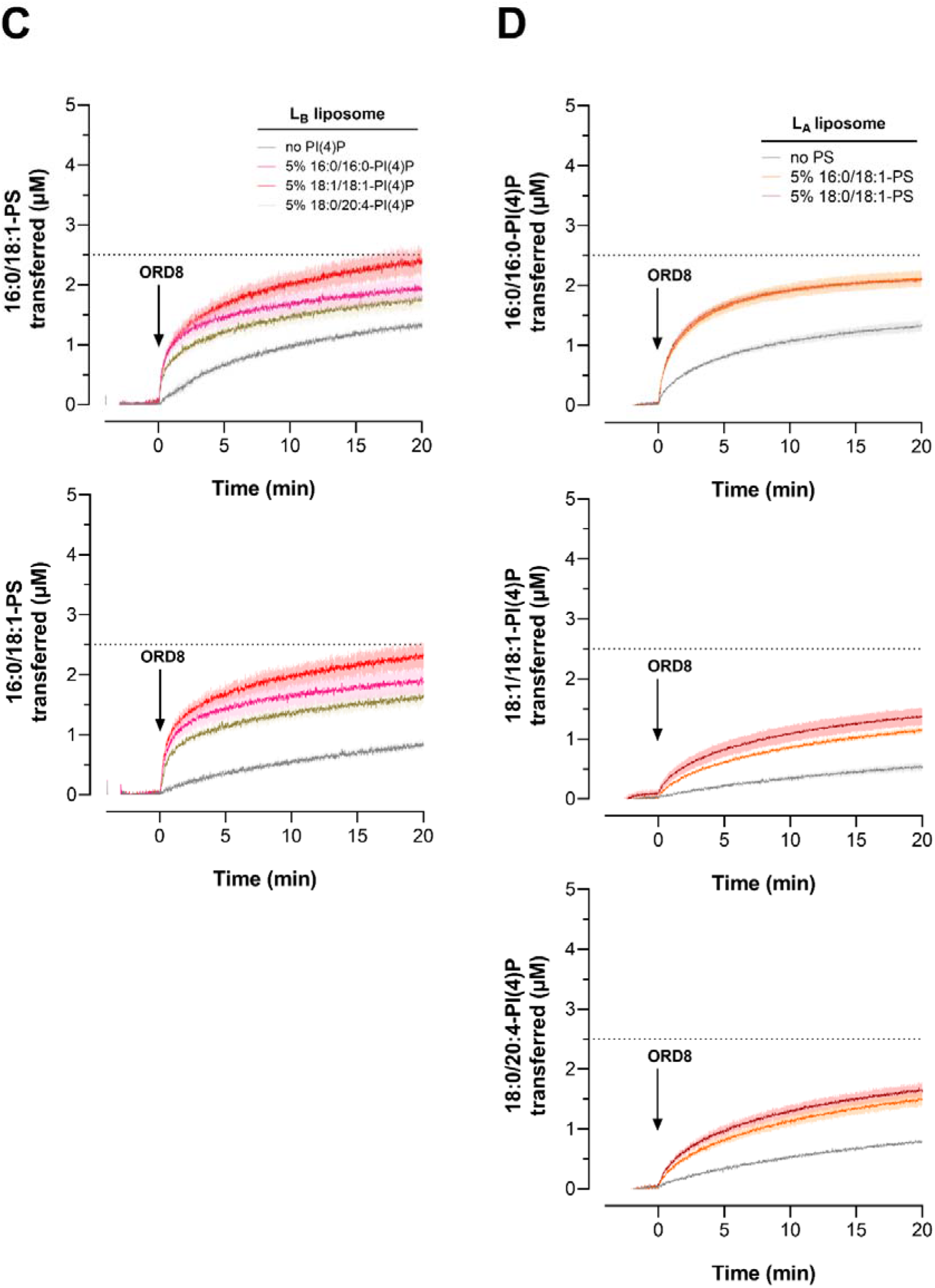
Transfer kinetics of 16:0/18:1-PS and 18:0/18:1-PS and PI(4)P with different unsaturation degree, measured with Osh6p or ORD8 in non-exchange and exchange conditions with different PS and PI(4)P combinations. (**A**) DOPC liposomes (200 µM total lipid, L_A_) containing 2% Rhod-PE and 5% 16:0/18:1-PS or 18:0/18:1-PS were mixed with NBD-C2_Lact_. After one minute, DOPC liposomes (200 µM lipids, L_B_) containing or not 5% of a particular PI(4)P species were added. After 3 min, Osh6p was injected. **(B)** L_B_ liposomes (200 µM lipids) containing a given PI(4)P species (at 5%) were pre-incubated with NBD-PH_FAPP_. Next, L_A_ liposomes (200 µM total lipid) containing 2% Rhod-PE, enriched or not with 5% 16:0/18:1-PS or 18:0/18:1-PS, and Osh6p were sequentially added. Experiences were conducted at 30°C with 200 nM Osh6p.**(C)** Measurements were taken as in **(A)** but with 240 nM ORD8 at 37°C and using 18:1/18:1-PI(4)P instead of 16:0/18:1-PI(4)P. **(D)** Same experiments as in **(B)** but performed with ORD8 and by replacing 16:0/18:1-PI(4)P by 18:1/18:1-PI(4)P. Each trace is the mean ± s.e.m. of independent kinetics (n = 3-4).

**Figure 4-Figure Supplement 1.**
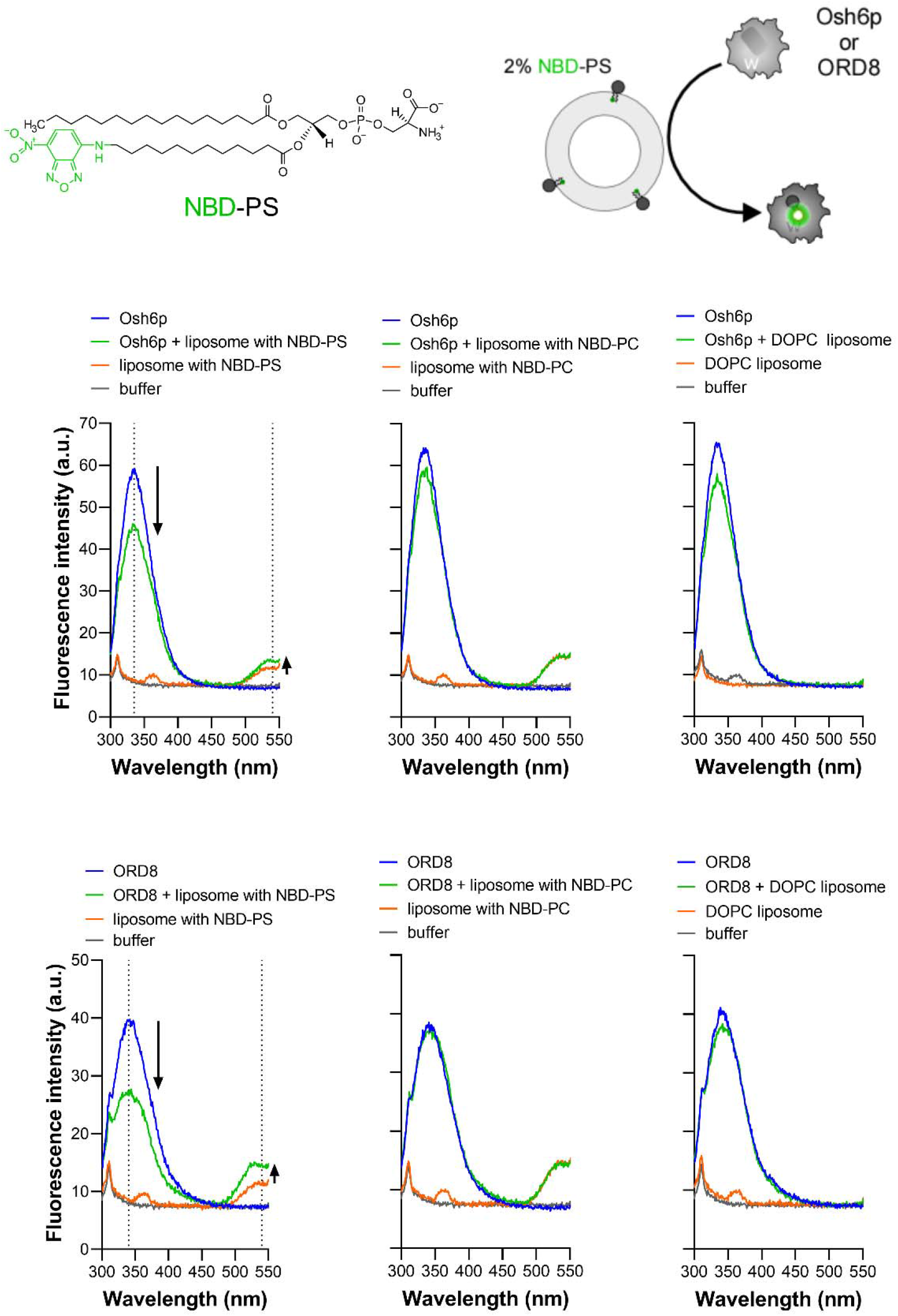
FRET between Osh6p or ORD8 and NBD-PS. Emission spectra of Osh6p and ORD8 (240 nM) alone in HK buffer or in the presence of liposomes (100 µM total lipid) containing either 2% NBD-PS or NBD-PC, measured at 30°C upon excitation at 280 nm. A control experiment was done with pure DOPC liposomes. For each condition, a spectrum was recorded in the absence of protein. Arrows indicate the diminution of the Trp fluorescence and the increase in NBD fluorescence that were observed once liposomes doped with NBD-PS were added to Osh6p or ORD8, indicative of a FRET process between each protein and the fluorescent lipid.

**Figure 4-Figure Supplement 2.**
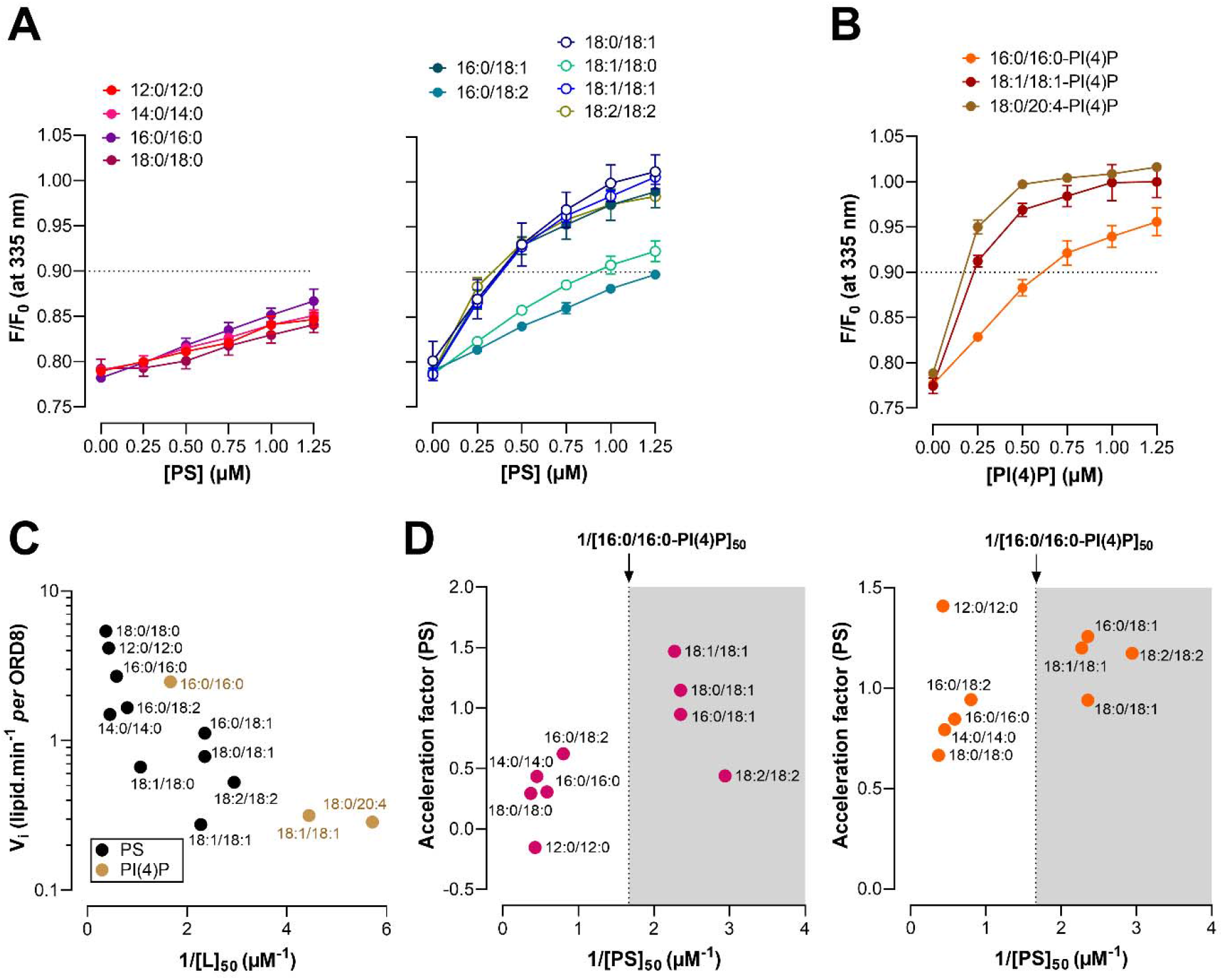
Relationship between the affinity of ORD8 for PS and PI(4)P species and its ability to transfer them. **(A)** Competition assays. DOPC liposomes (100 µM total lipid, final concentration) doped with 2% NBD-PS, were added to ORD8 (240 nM) in HK buffer at 30°C. The sample was excited at 280 nm and the emission was measured at 335 nm. Incremental amounts of liposome, containing a given PS species at 5%, were injected to the sample. The fluorescence was normalized considering the initial F_max_ fluorescence, prior to the addition of NBD-PS-containing liposomes, and the dilution effect due to liposomes addition. Data are represented as mean ± s.e.m. (n = 3). **(B)** Same experiments as in (**A)** except that liposomes containing a given PI(4)P species at 5% were injected. Data are represented as mean ± s.e.m. (n = 4 with 16:0/16:0-PI(4)P, n = 3 for other PI(4)P species). **(C)** Initial transfer rates determined under non-exchange conditions for PS and PI(4)P subspecies with ORD8 (shown in Figure 1c, d and 2a) as a function of 1/[L]_50_ values (right panel) determined for each lipid subspecies. **(D)** Acceleration factors determined from PS and 16:0/16:0-PI(4)P transfer rates obtained in the experiments shown in Figure 1B and C, as a function of the 1/[L]_50_ value determined for each PS subspecies.

**Figure 5-Figure Supplement 1.**
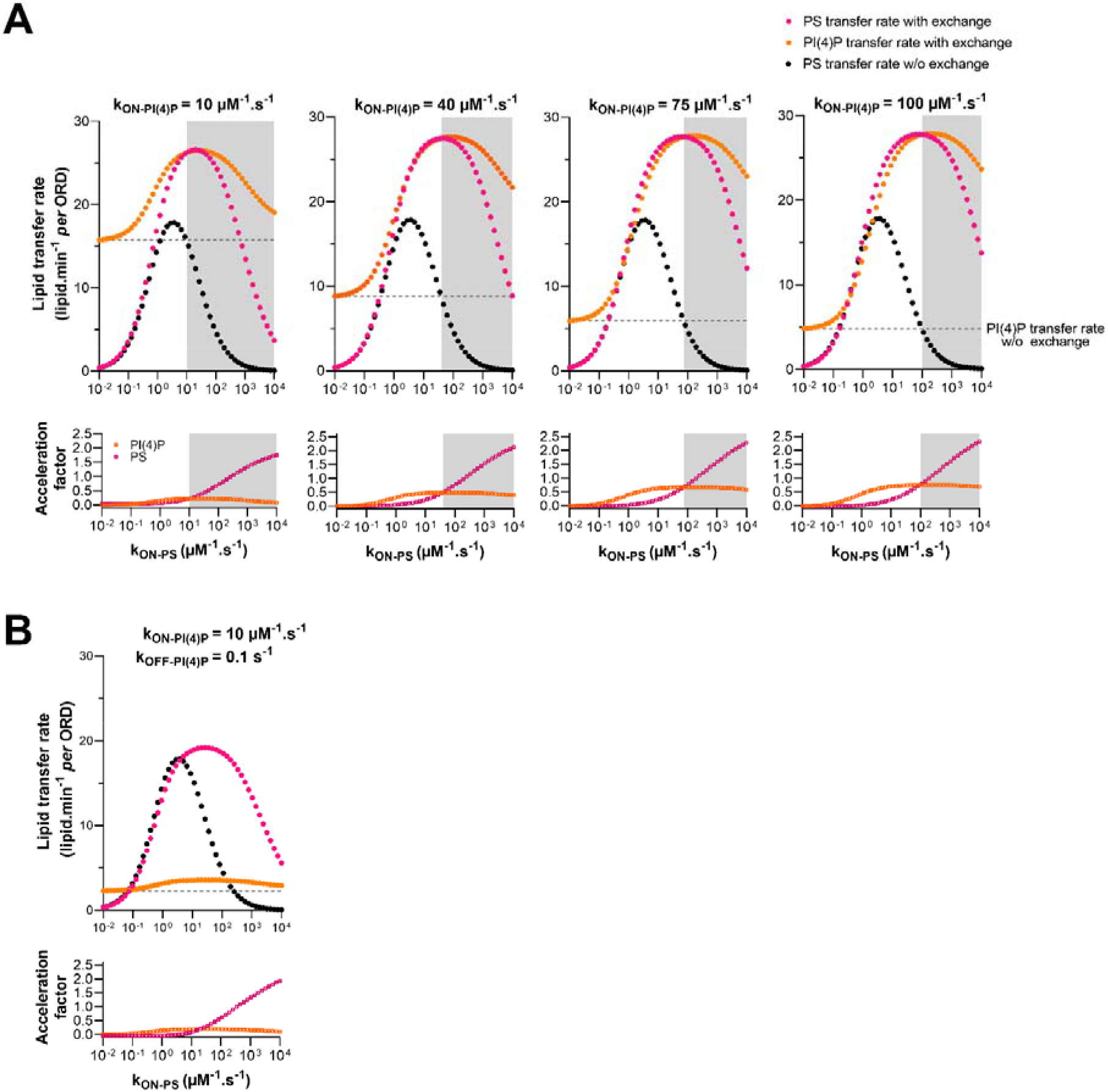
Analysis of the relation between ORD’s affinity for PS and PI(4)P and its ability to transfer these lipids between membranes. **(A)** Initial PS (pink dots) and PI(4)P transfer rates (orange dots) were calculated as a function of k_ON-PS_ for different k_ON_-_PI(4)P_ values (10 µM^−1^.s^−1^, as shown in Figure 5, but also 40, 75 and 100 µM^−1^.s^−1^), considering that PS and PI(4)P were initially present at 5% in the A and B membranes, respectively (exchange condition). The PS transfer rates, calculated as a function of k_ON-PS_ values, and the PI(4)P transfer rate under non-exchange condition are shown (black dots and dashed line, respectively). The grey areas correspond to regimes where k_ON-PS_ > k_ON-PI(4)P_ *i.e.,* the ORD has more affinity for PS than PI(4)P. The acceleration factors, calculated for PS and PI(4)P are shown. **(B)** Simulations were carried out as in **(A)** with k_ON-PI(4)P_ = 10 µM^−1^.s^−1^ and k_OFF-PI(4)P_ = 0.1 s^−1^. Acceleration factors are shown.

**Figure 6-Figure Supplement 1.**
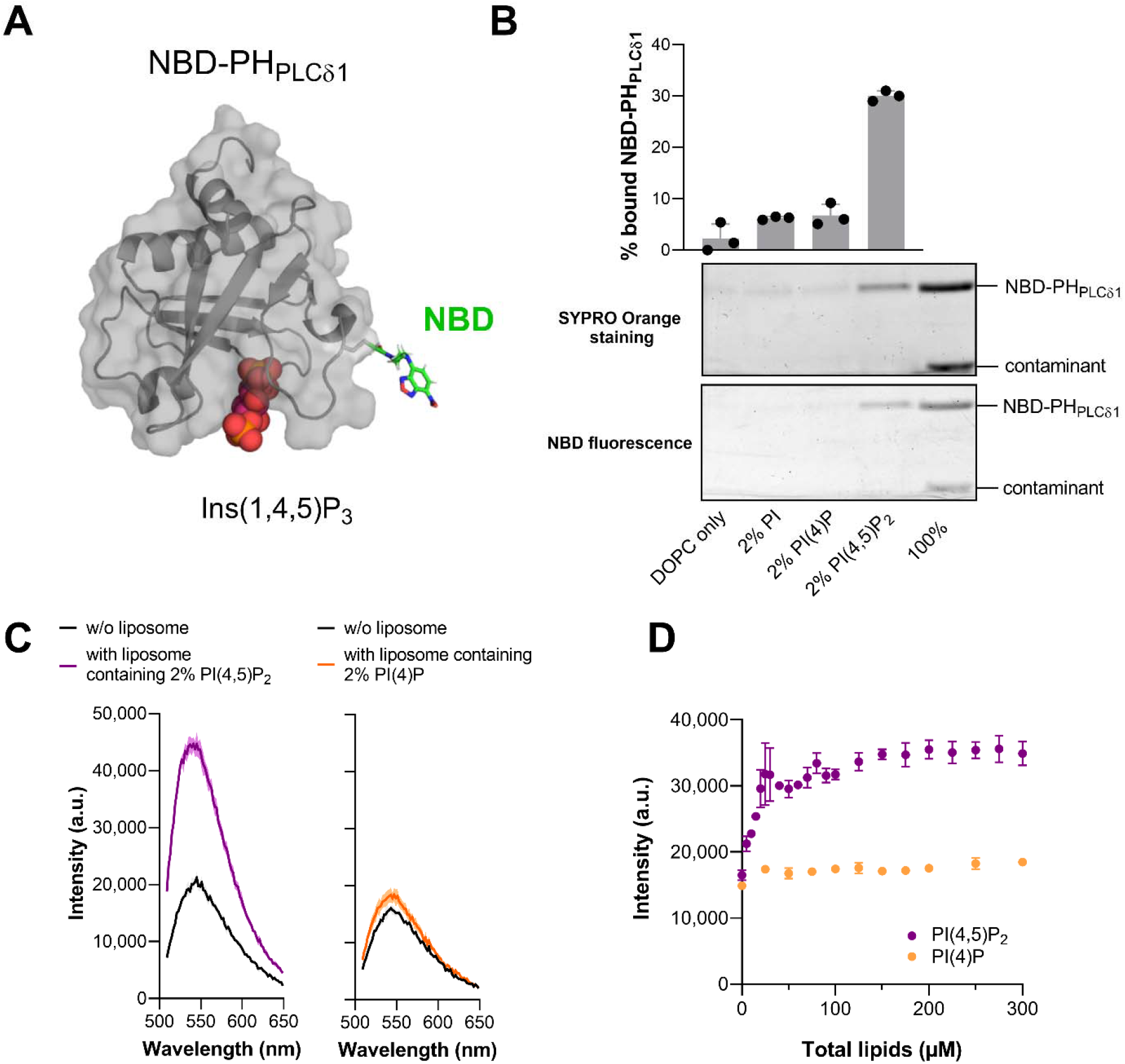
Biochemical characterization of NBD-PH_PLCδ1_. **(A)** Tridimensional model of the NBD-labeled PH_PLCδ1_ based on the crystal structure of the PH domain of the *Rattus norvegicus* Phospholipase C δ1 (PDB ID: 1MAI^43^). The solvent-exposed cysteine C48 was mutated into a serine; serine S61 was replaced by a cysteine. An N,N’-dimethyl-N-(thioacetyl)-N’-(7-nitrobenz-2-oxa-1,3-diazol-4-yl)ethylenediamine moiety (in stick, with carbon in green, nitrogen in blue and oxygen in red), built manually and energetically minimized, is grafted to the thiol function of C61. The inositol 1,4,5-trisphosphate (Ins(1,4,5)P_3_) molecule, corresponding to the polar head of PI(4,5)P_2_ is represented in sphere mode with oxygen in red and phosphate in orange. The figure was prepared with PyMOL (http://pymol.org/). **(B)** Flotation assays. NBD-PH_PLCδ1_ (1 µM) was incubated in HK buffer at room temperature for 10 min under agitation with liposomes (1.5 mM total lipid concentration) only made of DOPC or containing 2% liver PI, 2% 18:0/20:4-PI(4)P or 2% 18:0/20:4-PI(4,5)P_2_. After centrifugation, the liposomes were recovered at the top of sucrose cushions and analyzed by SDS-PAGE. The presence of NBD-PH_PLCδ1_ in the top fraction (lane 1 to 4) and in the reference lane (lane 5) was visualized by direct fluorescence, and after staining the gel with SYPRO Orange, using a fluorescence imaging system. The amount of protein recovered in the top fraction was quantified based on the SYPRO Orange signal and the percentage of NBD-PH_PLCδ1_ bound to liposomes was determined using the content of lane 5 (100%) as a reference. Data are represented as mean ± s.e.m. (n = 3). **(C)** Fluorescence spectra of NBD-PH_PLCδ1_ (250 nM) in buffer alone or mixed with DOPC liposome (350 µM) containing either 2% PI(4)P or 2% PI(4,5)P_2_. **(D)** NBD intensity measured at 535 nm as function of total lipid concentration and with liposome composed of DOPC, DOPC/PI (98:2), DOPC/PI(4)P (98:2) or DOPC/PI(4,5)P_2_ (98:2). Data are represented as mean ± s.e.m. (n = 3).

**Figure 6-Figure Supplement 2.**
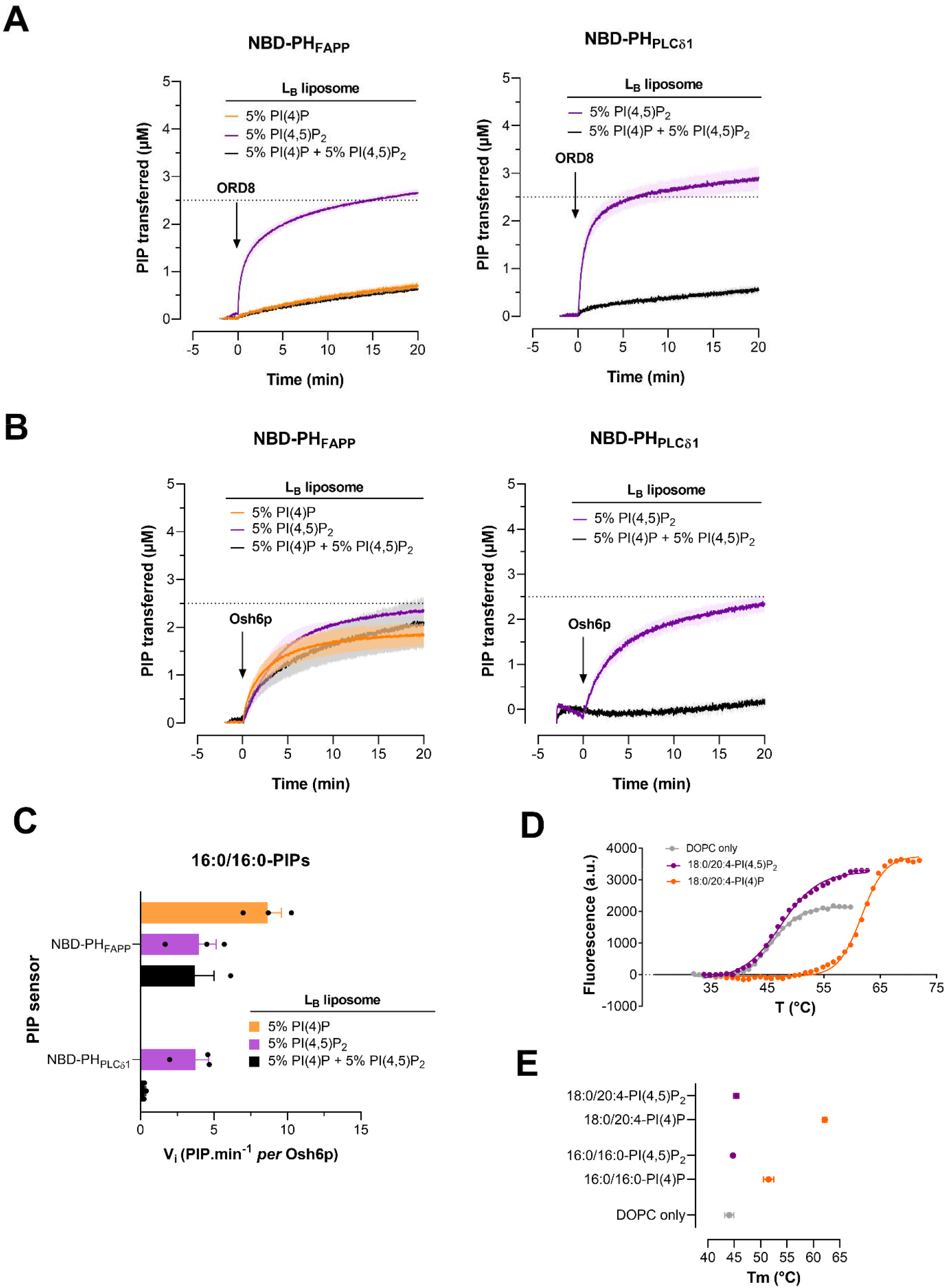
PIP selectivity of ORP8 and Osh6p. **(A)** Intermembrane PI(4)P and PI(4,5)P_2_ transfer activity of ORD8 or Osh6p when PI(4)P, PI(4,5)P_2_ or both PIPs were present in donor L_B_ liposomes. Experiments were performed with ORD8 as for those shown in Figure 6A except that 18:0/20:4-PIPs were assayed. The injection of the LTP set the time to zero. Each trace is the mean ± s.e.m. of independent kinetics (n = 3). **(B)** Intermembrane PIP transfer activity of Osh6p. Experiments were performed at 30°C as described in Figure 6A using Osh6p (200 nM final concentration) and liposomes containing PI(4)P and PI(4,5)P_2_ with 16:0/16:0 acyl chains. Each trace is the mean ± s.e.m. of independent kinetics (n = 3). **(C)** Initial transfer rates of 16:0/16:0-PIPs determined with NBD-PH_FAPP_ and NBD-PH_PLCδ1_ based on the kinetic curves measured with Osh6p and shown in panel (**B)**. Data are represented as mean ± s.e.m. (error bars, n = 3). **(D)** Melting curves of Osh6p loaded with 18:0/20:4-PI(4)P or 18:0/20:4-PI(4,5)P_2_. A control experiment with Osh6p incubated with DOPC liposomes without ligands is shown. **(E)** Melting temperatures□(T_m_) determined for Osh6p incubated with liposome devoid of lipid ligand (DOPC only) or loaded with PI(4)P or PI(4,5)P_2_ with 16:0/16:0 or 18:0/20:4 composition. Data are represented as mean ± s.e.m (n = 3) except for 18:0/20:4-PI(4)P and 16:0/16:0-PI(4,5)P_2_ (n = 2).

**Figure 7-Figure Supplement 1.**
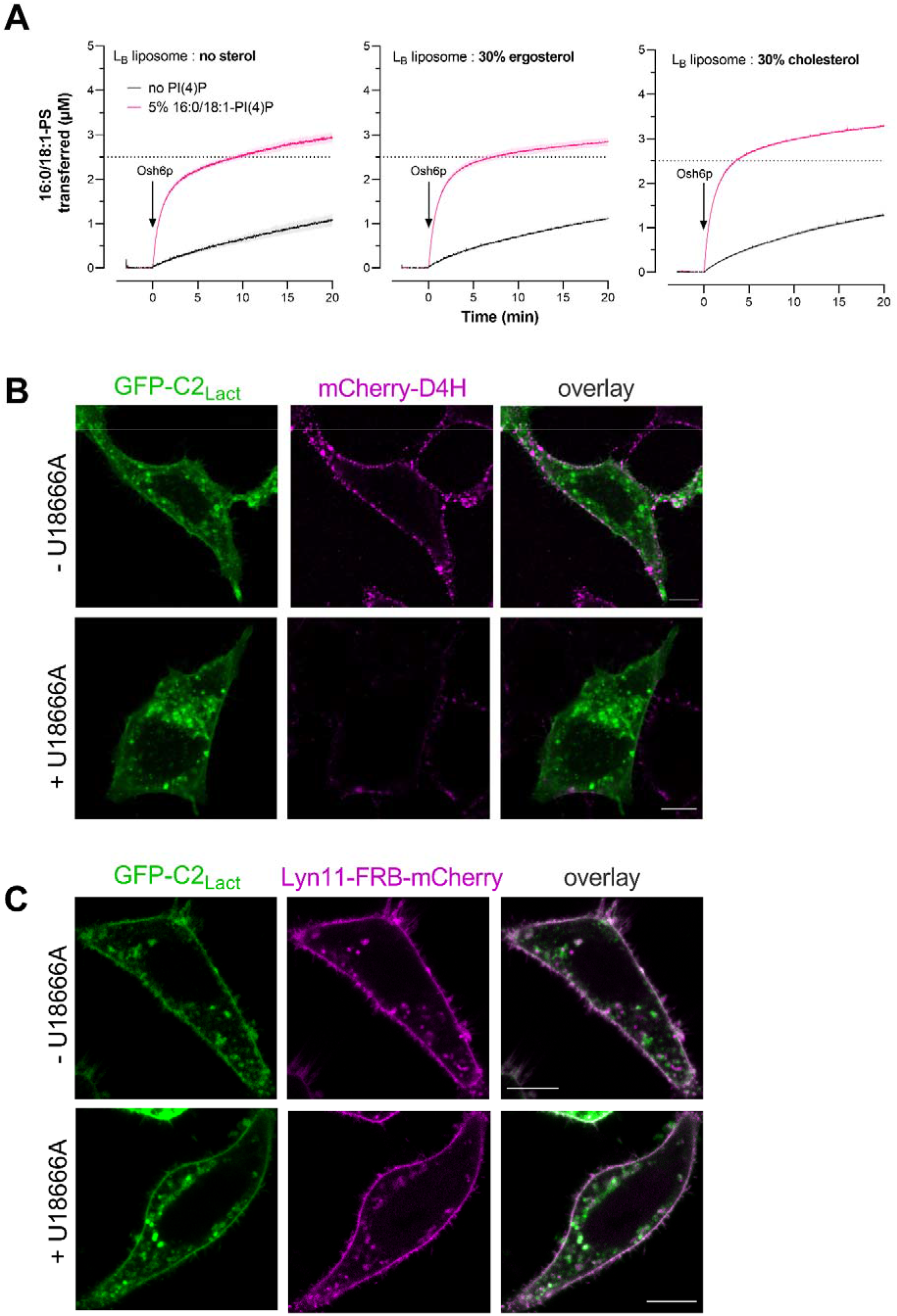
Impact of sterol on PS/PI(4)P exchange. **(A)** Transfer kinetics of 16:0/18:1-PS from L_A_ liposomes (200 µM total lipid, 5% PS) to L_B_ liposomes (200 µM) without or with 30% sterol (cholesterol or ergosterol), doped or not with 5% 16:0/18:1-PI(4)P (at the expense of PC). Measurements were done at 30°C with Osh6p (200 nM). Each trace is the mean ± s.e.m. of kinetics recorded in independent experiments (n = 4-5). **(B)** Confocal images of live GFP□C2_Lact_□expressing HeLa cells (green) treated or not with U18666A for 24 h at 37°C. Cholesterol in the PM was detected by incubating the cells for 10 min with mCherry-D4H (purple) at room temperature and then washed with medium prior to imaging. The overlay panel shows merged green and magenta images. Scale bars: 10 µm. **(C)** Confocal images of live cells that co-express GFP□C2_Lact_ (green) and Lyn11-FRB-mCherry (magenta) and treated or not with U18666A for 24 h at 37°C. The treatment with the drug did not change the localization of Lyn11-FRB-mCherry used as a stable marker of the PM. The overlay panel shows merged green and magenta images. Scale bars: 10 µm.

## Material and Methods

### Protein expression, labelling and purification

Osh6p, Osh6p(noC/S190C), NBD-PH_FAPP_ and NBD-C2_Lact_ were purified as previously described^13,44,58^. Their concentration was determined by UV spectrometry.

GST-ORD8 (GST-ORP8[370-809])^14^ was expressed in *E. Coli* (BL21-GOLD(DE3)) competent cells (Stratagene) grown in Luria Bertani Broth (LB) medium at 18°C overnight upon induction with 0.1 mM isopropyl β-D-1-thiogalactopyranoside (IPTG). When the optical density of the bacterial suspension, measured at 600 nm (OD_600_), reached a value of 0.6, bacteria cells were harvested and re-suspended in cold buffer (50 mM Tris, pH 8, 500 mM NaCl, 2 mM DTT) supplemented with 1 mM PMSF, 10 µM bestatin, 1 µM pepstatin A and cOmplete EDTA-free protease inhibitor tablets (Roche). Cells were lysed in a Cell Disruptor TS SERIES (Constant Systems Ltd.) and the lysate was centrifuged at 186,000 × g for 1 h. Then, the supernatant was applied to Glutathione Sepharose 4B (Cytiva) for 4 h at 4°C. After three washing steps with the buffer devoid of protease inhibitors, the beads were incubated with PreScission Protease (Cytiva) overnight at 4°C to cleave off the ORD8 from the GST domain. The protein was recovered in the supernatant after several cycles of centrifugation and washing of the beads, concentrated and injected onto a XK-16/70 column packed with Sephacryl S-200 HR to be purified by size-exclusion chromatography. The fractions with ∼100% pure ORD8 were pooled, concentrated and supplemented with 10% (v/v) pure glycerol (Sigma). Aliquots were prepared, flash-frozen in liquid nitrogen and stored at −80°C. The concentration of the protein was determined by measuring its absorbance at λ = 280 nm (ε = 81,820 M^−1^.cm^−1^).

To prepare NBD-labelled PH_PLCδ1_, an endogenous, solvent-exposed cysteine at position 48 of a GST-PH_PLCδ1_ construct^59^ (PH domain of 1-phosphatidylinositol 4,5-bisphosphate phosphodiesterase delta-1, *R. norvegicus*, Uniprot: P10688) was replaced by a serine, and a serine at position 61 was replaced by a cysteine using the Quikchange kit (Agilent). GST-PH_PLCδ1_ was expressed in *E. Coli* BL21-GOLD(DE3) competent cells at 20°C for 24 h upon induction with 0.1 mM IPTG (at OD_600_ = 0.6). Harvested bacteria cells were re-suspended in 50 mM Tris, pH 7.4, 120 mM NaCl buffer containing 2 mM DTT and supplemented with 1 mM PMSF, 10 µM bestatin, 1 µM pepstatin A and EDTA-free protease inhibitor tablets. Cells were lysed and the lysate was centrifuged at 186,000 × g for 1 h. Next, the supernatant was applied to Glutathione Sepharose 4B for 4 h at 4°C. The beads were washed three times with protease inhibitor-free buffer and incubated with thrombin at 4°C for 16 h to cleave off the protein from the GST domain. Then, the eluate obtained after thrombin treatment was concentrated and mixed (after DTT removal by gel filtration on illustra NAP-10 columns (Cytivia) with a 10-fold excess of N,N’-dimethyl-N-(iodoacetyl)-N’-(7-nitrobenz-2-oxa-1,3-diazol-4-yl) ethylenediamine (IANBD-amide, Molecular Probes). After 90 min on ice, the reaction was stopped by adding a 10-fold excess of L-cysteine over the probe. The free probe was removed by size-exclusion chromatography using a XK-16/70 column packed with Sephacryl S-200 HR. The fractions that contained NBD-PH_PLCδ1_ were pooled, concentrated, supplemented with 10% (v/v) pure glycerol. Aliquots were stored at −80°C once flash-frozen in liquid nitrogen. The labelled protein was analyzed by SDS-PAGE and UV-visible spectroscopy. The gel was directly visualized in a fluorescence imaging system (FUSION FX fluorescence imaging system) to detect NBD-labelled protein excited in near-UV and then stained with SYPRO Orange to determine the purity of NBD-labelled protein. The labelling yield (∼100%) was estimated from the ratio between the absorbance of the protein at λ = 280 nm (ε = 17,990 M^−1^.cm^−1^ for PH_PLCδ1_) and NBD at λ = 495 nm (ε = 25,000 M^−1^ cm^−1^). The concentration of the protein was determined by a BCA assay and UV-spectrometry.

mCherry-D4-His_6_ and mCherry-D4H-His_6_ (carrying the D434S mutation) were each overexpressed overnight in *E. Coli* (BL21-GOLD(DE3)) at 18°C for 20 h upon induction by 0.4 mM IPTG at OD_600_ = 0.6. Bacteria cells were harvested and re-suspended in 50 mM NaH_2_PO_4_/Na_2_HPO_4_, pH 8, 300 mM NaCl, 10 mM imidazole buffer supplemented with 1 mM PMSF, 10 µM bestatine, 1 µM pepstatine and EDTA-free protease inhibitor tablets. Cells were broken and the lysate was centrifuged at 186,000 × g for 1 h. Then, the supernatant was applied to HisPur Cobalt Resin (Thermo Scientific) for 4 h at 4°C. The beads were loaded into a column and washed four times with buffer devoid of protease inhibitors. Bound protein was eluted from the beads incubated for 10 minutes with 20 mM NaH_2_PO_4_/Na_2_HPO_4_, pH 7.4, 250 mM imidazole buffer. This step was repeated six times to collect a maximal amount of protein. Each protein was concentrated and stored at −80°C in the presence of 10% (v/v) glycerol. The concentration of protein was determined by measuring the absorbance at λ = 280 nm (ε = 77,810 M^−1^ .cm^−1^).

### Lipids

18:1/18:1-PC (1,2-dioleoyl-*sn*-glycero-3-phosphocholine or DOPC), 12:0/12:0-PS (1,2-dilauroyl-*sn*-glycero-3-phospho-L-serine or DLPS), 14:0/14:0-PS (1,2-dimyristoyl-*sn*-glycero-3-phospho-L-serine or DMPS), 16:0/16:0-PS (1,2-dipalmitoyl-*sn*-glycero-3-phospho-L-serine or DPPS), 18:0/18:0-PS (1,2-distearoyl-*sn*-glycero-3-phospho-L-serine or DSPS), 16:0/18:1-PS (1-palmitoyl-2-oleoyl-*sn*-glycero-3 phospho-L-serine or POPC), 18:0/18:1-PS (1-stearoyl-2-oleoyl-*sn*-glycero-3-phospho-L-serine or SOPS), 18:1/18:0-PS (1-oleoyl-2-stearoyl-*sn*-glycero-3-phospho-L-serine or OSPS), 18:1/18:1-PS (1,2-dioleoyl-*sn*-glycero-3-phospho-L-serine or DOPS), 16:0/18:2-PS (1-palmitoyl-2-linoleoyl-*sn*-glycero-3-phospho-L-serine), 18:2/18:2-PS (1,2-dilinoleoyl-*sn*-glycero-3-phospho-L-serine), liver PI (L-α-phosphatidylinositol, bovine), 16:0/18:1-PI(4)P (1-palmitoyl-2-oleoyl-*sn*-glycero-3-phospho-(1’-myo-inositol-4’-phosphate)), 18:1/18:1-PI(4)P (1,2-dioleoyl-*sn*-glycero-3-phospho-(1’-myo-inositol-4’-phosphate)), brain PI(4)P (L-α-phosphatidylinositol 4-phosphate), brain PI(4,5)P_2_ (L-α-phosphatidylinositol 4,5-bisphosphate), Rhodamine-PE (1,2-dipalmitoyl-*sn*-glycero-3-phosphoethanolamine-N-(lissamine rhodamine B sulfonyl)), 16:0/12:0 NBD-PS (1-palmitoyl-2-(12-[(7-nitro-2-1,3-benzoxadiazol-4-yl)amino]dodecanoyl)-*sn*-glycero-3-phosphoserine), 16:0/12:0 NBD-PC (1-palmitoyl-2-(12-[(7-nitro-2-1,3-benzoxadiazol-4-yl)amino]dodecanoyl)-*sn*-glycero-3-phosphocholine), cholesterol and ergosterol were purchased from Avanti Polar Lipids. Saturated PIPs, namely 16:0/16:0-PI(4)P (1,2-dipalmitoyl-*sn*-glycero-3-phospho-(1’-myo-inositol-4’-phosphate) and 16:0/16:0-PI(4,5)P_2_ (1,2-dipalmitoyl-*sn*-glycero-3-phospho-(1′-myo-inositol-4′,5′-bisphosphate)) were purchased from Echelon Biosciences.

### Liposomes preparation

Lipids stored in stock solutions in CHCl_3_ or CHCl_3_/methanol were mixed at the desired molar ratio. The solvent was removed in a rotary evaporator under vacuum. If the flask contained a mixture with PI(4)P and/or PI(4,5)P_2_, it was pre-warmed at 33°C for 5 min prior to creating a vacuum. The lipid film was hydrated in 50 mM HEPES, pH 7.2, 120 mM K-Acetate (HK) buffer to obtain a suspension of multi-lamellar vesicles. The multi-lamellar vesicles suspensions were frozen and thawed five times and then extruded through polycarbonate filters of 0.2 µm pore size using a mini-extruder (Avanti Polar Lipids). Liposomes were stored at 4°C and in the dark when containing fluorescent lipids and used within 2 days.

### Lipid transfer assays with two liposome populations

Lipid transfer assays were carried out in a Shimadzu RF 5301-PC or a JASCO FP-8300 spectrofluorometer. Each sample (600 µL final volume) was placed in a cylindrical quartz cuvette, continuously stirred with a small magnetic bar and thermostated at 30°C and 37°C for experiments done with Osh6p and ORD8, respectively. At precise times, samples were injected from stock solutions with Hamilton syringes through a guide in the cover of the spectrofluorometer. The signal of fluorescent lipid sensors (NBD-C2_Lact_, NBD-PH_FAPP_ and NBD-PH_PLCδ_) was followed by measuring the NBD signal at λ = 530 nm (bandwidth 10 nm) upon excitation at λ = 460 nm (bandwidth 1.5 nm) with a time resolution of 0.5 s. To measure PS transfer, a suspension (540 µL) of L_A_ liposomes (200 µM total lipid, final concentration), made of DOPC and containing 5% PS and 2% Rhod-PE, was mixed with 250 nM NBD-C2_Lact_ in HK buffer. After 1 min, 30 µL of a suspension of L_B_ liposomes (200 µM lipids, final concentration) containing or not 5% PI(4)P was added to the sample. Three minutes after, Osh6p (200 nM) or ORD8 (240 nM) was injected. The amount of PS ([PS], expressed in µM) transferred over time was determined from the raw NBD traces using the formula [PS] = 2.5 × F_Norm_ with F_Norm_ = (F-F_0_)/(F_Eq_-F_0-Eq_). F corresponds to the data point recorded over time. F_0_ is equal to the average NBD signal measured between the injection of L_B_ liposomes and LTP. F_Eq_ is the signal measured with the sensor in the presence of L_A-Eq_ and L_B-Eq_ liposomes, and F_0-Eq_ is the signal of the suspension of L_A-Eq_ only, prior to the addition of L_B_-_Eq_. The lipid composition of L_A-Eq_ and L_B-Eq_ liposomes were similar to that of L_A_ and L_B_ liposomes used in the transfer assays, except that both contained 2.5% PS (and additionally 2.5% PI(4)P in the experiments conducted in the context of the lipid exchange) to normalize the signal. The amount of PS transferred from L_A_ to L_B_ liposomes corresponds to 2.5 × F_Norm_, as one considers that at equilibrium one half of accessible PS molecules, contained in the outer leaflet of the L_A_ liposomes *(i.e*., corresponding to 5% of 0.5 × 200 µM total lipids) have been delivered into L_B_ liposomes.

To measure PI(4)P transfer, a suspension (540 µL) of L_B_ liposomes (200 µM total lipid) containing 5% PI(4)P was mixed with 250 nM NBD-PH_FAPP_ in HK buffer. After 1 minute, 30 µL of a suspension of L_A_ liposomes (200 µM lipids) containing 2% Rhod-PE and doped or not with 5% PS were injected. After 3 additional minutes, LTP was injected. The amount of PI(4)P transferred from L_B_ to L_A_ liposomes was determined considering that [PI(4)P] = 2.5 × F_Norm_ with F_Norm_ = (F-F_0_)/(F_Eq_-F_0_). F corresponded to the data point recorded over time, F_0_ was the average signal measured before the addition of LTP and F_Eq_ was the average signal measured in the presence of L_A-Eq_ and L_B-Eq_ liposomes that each contained 2.5% PI(4)P (and additionally 2.5% PS to normalize data obtained in exchange conditions). At equilibrium, it is considered that one half of accessible PI(4)P molecules, contained in the outer leaflet of L_B_ liposomes, (*i.e.,* corresponding to 5% of 0.5 × 200 µM total lipids) have been transferred into L_A_ liposomes. The transfer of PI(4,5)P_2_ from L_B_ to L_A_ liposomes was determined using NBD-PH_FAPP_ or NBD-PH_PLCδ1_ at 250 nM as described for PI(4)P transfer measurements except that F_Eq_ was determined using L_A_ and L_B_ liposomes that each contained 2.5% PI(4,5)P_2_. For all the measurements, the initial transport rates (or initial velocities) were determined from normalized curves by fitting the first eight data points (4 s) measured upon Osh6p/ORD8 injection with a linear function, divided by the amount of LTP and expressed in terms of lipid.min^−1^per protein.

### PS transport assay with three liposome populations

At time zero, L_A_ liposomes (200 µM total lipid, final concentration) containing 95% DOPC and 5% 16:0/18:1-PS were mixed with 250 nM NBD-C2_Lact_ in 480 µL of HK buffer at 37°C in a quartz cuvette. After 1 min, 60 µL of a suspension of L_B_ liposomes (200 µM total lipid) made of 93% DOPC, 2% Rhod-PE and 5% 18:0/20:4-PI(4)P or 18:0/20:4-PI(4,5)P_2_ were injected. After 2 additional minutes, 60 µL of a suspension of L_C_ liposomes (200 µM total lipid) consisting only of DOPC or containing 5% 18:0/20:4-PI(4)P or 18:0/20:4-PI(4,5)P_2_. Finally, after 2 minutes, ORD8 (240 nM) was injected. PS transport was followed by measuring the NBD signal at λ = 525 nm (bandwidth 5 nm) upon excitation at λ = 460 nm (bandwidth 1 nm) under constant stirring. The quenching of the NBD signal was due to the translocation of the probe onto L_B_ liposome doped with Rhod-PE, and reflected how much PS was transferred from L_A_ to L_B_ liposomes. The amount of transferred PS (in µM) was determined by normalizing the NBD signal considering that [PS] = 2.5 × F_Norm_ with F_Norm_ = (F-F_0_)/(F_Eq_-F_0_). F corresponded to data points measured over time, F_0_ corresponded to the NBD-C2_Lact_ signal measured when PS was only in L_A_ liposomes and F_Eq_ corresponded to the signal of the probe when PS was fully equilibrated between L_A_ and L_B_ liposomes. F_0_ value was obtained by averaging the fluorescence measured after the injection of L_C_ liposomes and before the injection of ORD8. F_Eq_ was the fluorescence of NBD-C2_Lact_ (250 nM) measured once it was sequentially mixed with L_A-Eq_ liposomes consisting of 97.5% DOPC and 2.5% 16:0/18:1-PS and L_B-Eq_ liposomes consisting of 95.5% DOPC, 2.5% 16:0/18:1-PS and 2% Rhod-PE, and L_C_ liposomes only made of DOPC (the concentration of each liposome population was 200 µM). Its value corresponded to the average fluorescence measured 15 min after the addition of L_C_ liposomes and for a 5-min period. F_Eq_ was also measured using only L_A_ and L_B_ liposomes and was found to be identical. A maximum of 2.5 µM of PS can be transferred from L_A_ to L_B_ liposomes as one half of accessible PS molecules, contained in the outer leaflet of the L_A_ liposomes, (*i.e.,* 5% of 0.5 × 200 µM total lipids) can be transferred to reach equilibrium.

### NBD-PS based competition assays

In a cylindrical quartz cuvette, Osh6p or ORD8 was diluted at 240 nM in a final volume of 555 µL of freshly degassed and filtered HK buffer at 30°C under constant stirring. Two minutes after, 30 µL of a suspension of DOPC liposomes containing 2% NBD-PS was added (100 µM total lipid, 1 µM accessible NBD-PS). Five minutes after, successive injections of 3 µL of a suspension of DOPC liposomes enriched with a given PS or PI(4)P species (at 5%) were done every 3 minutes. Tryptophan fluorescence was measured at λ = 340 nm (bandwidth 5 nm) upon excitation at λ = 280 nm (bandwidth 1.5 nm). The signal was normalized by dividing F, the signal measured over time, by F_0_, the signal measured prior to the addition of the NBD-PS-containing liposome population, and corrected for dilution effects due to the successive injections of the second population of liposome. The signal between each liposome injection was averaged over 2 min to build the binding curve as a function of concentration of accessible non-fluorescent PS or PI(4)P species (from 0 to 1.25 µM).

### Thermal shift assay

The relative melting temperatures (T_m_) of Osh6p in an empty form or loaded with a lipid ligand were determined by measuring the unfolding of the protein as a function of increasing temperature through the detection of the denatured form of the protein by fluorescence. To prepare Osh6p-PS and Osh6p-PI(4)P complexes, the protein at 5 µM was incubated with heavy liposomes made of DOPC (800 µM total lipid), containing a given PS or PI(4)P subspecies (5%) and encapsulating 50 mM HEPES, pH 7.4, 210 mM sucrose buffer, in a volume of 250 µL of HK buffer. An apo form of the protein was prepared by incubating the protein with DOPC liposome devoid of lipid ligands. Each sample was mixed by agitation for 30 min at 30°C and then were centrifuged at 400,000 × g for 20 min at 20°C to pellet the liposomes using a fixed-angle rotor (Beckmann TLA 120.1). A fraction of each supernatant (200 µL) containing Osh6p loaded with lipid was collected and the concentration of each complex was assessed by measuring sample absorbance. A volume of 15 µL of each Osh6p sample was mixed with 5× SYPRO Orange in an individual well of a 96-well PCR plate. The plates were sealed with an optical sealing tape (Bio-Rad) and heated in an Mx3005P Q-PCR system (Stratagene) from 25 to 95°C with a step interval of 1°C. The excitation and emission wavelengths were set at λ = 545 nm and λ = 568 nm, respectively (Cy3 signal). Fluorescence changes in the wells were measured with a photomultiplier tube. The melting temperatures (T_m_) were obtained by fitting the fluorescence data from 3-5 independent experiments with a Boltzmann model using the GraphPad Prism software.

### Flotation experiment

NBD-PH_PLCδ1_ protein (1 µM) was incubated with liposomes (1.5 mM total lipid) only made of DOPC, or additionally doped with 2% liver PI, 18:0/20:4-PI(4)P or 18:0/20:4-PI(4,5)P_2_, in 150 µL of HK buffer at room temperature for 10 min under agitation. The suspension was adjusted to 28% (w/w) sucrose by mixing 100 µL of a 60% (w/w) sucrose solution in HK buffer and overlaid with 200 µL of HK buffer containing 24% (w/w) sucrose and 50 µL of sucrose-free HK buffer. The sample was centrifuged at 240,000 × g in a swing rotor (TLS 55 Beckmann) for 1 h. The bottom (250 µL), middle (150 µL) and top (100 µL) fractions were collected. The bottom and top fractions were analyzed by SDS-PAGE by direct fluorescence and after staining with SYPRO Orange, using a FUSION FX fluorescence imaging system.

### Fluorescence-based membrane binding assay

Measurements were taken in a 96-well black plate (Microplate 96 Well PS F-Bottom Black Non-Binding, Greiner Bio-one) using a TECAN M1000 Pro. Incremental amounts of DOPC liposomes, containing either 2% 18:0/20:4-PI(4)P or 18:0/20:4-PI(4,5)P_2_, were mixed with NBD-PH_PLCδ1_ (250 nM) at 25°C in individual wells (100 µL final volume). NBD spectra were recorded from 509 to 649 nm (bandwidth 5 nm) upon excitation at 460 nm (bandwidth 5 nm). The intensity at λ = 535 nm was plotted as a function of total lipid concentration (from 0 to 300 µM).

### CPM accessibility assay

The day of the experiment, 100 µL from a stock solution of Osh6p(noC/S190C) construct was applied onto an illustra NAP□5 column (Cytivia) and eluted with freshly degassed HK buffer, according to the manufacturer’s indications to remove DTT. The concentration of the eluted protein was determined by UV□spectroscopy considering ε = 55,810 M^−1^.cm^−1^ at λ = 280 nm. A stock solution of CPM (7-Diethylamino-3-(4-maleimidophenyl)-4-methylcoumarin, Sigma-Aldrich) at 4 mg/mL was freshly prepared as described in ^60^ by mixing 1 mg of CPM powder in 250 µL of DMSO. Thereafter, this solution was diluted in a final volume of 10 mL of HK buffer and incubated for 5 min at room temperature. The solution was protected from light and used immediately. In individual wells of a 96-well black plate (Greiner Bio-one), Osh6p(noC/S190C) at 400 nM was mixed either with DOPC liposomes (400 µM total lipid) or liposomes containing 2% 18:0/20:4-PI(4)P or 18:0/20:4-PI(4,5)P_2_ in 200 µL of HK buffer. A small volume of CPM stock solution was then added to obtain a final concentration of 4 µM. After a 30 min-incubation at 30°C, emission fluorescence spectra were measured from 400 to 550 nm (bandwidth 5 nm) upon excitation at λ = 387 nm (bandwidth 5 nm) using a fluorescence plate reader (TECAN M1000 Pro). The maximal intensity of the spectral peak was at 460 nm. Control spectra were recorded in the absence of protein for each condition.

### Kinetic modeling

To analyze the experimental data, we considered that an ORD-mediated PS/PI(4)P exchange cycle can be described by the following sequence of reactions:

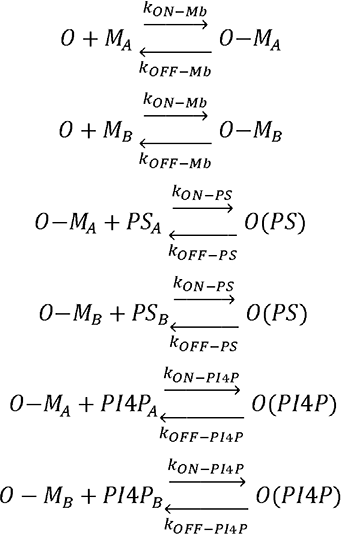

O corresponds to an empty form of the ORD in solution. O-M_A_ and O-M_B_ correspond to ORD in an empty state, bound to A and B membrane, respectively. O(PS) and O(PI4P) correspond to ORD in a soluble state in 1:1 complex with a PS and PI(4)P molecule, respectively. PS_A_ and PS_B_ correspond to the PS pool in the A and B membrane, respectively. PI4P_A_ and PI4P_B_ are the PI(4)P pools in the A and B membrane, respectively. The time evolution of the PS and PI(4)P concentrations in A and B membranes respectively was determined by integrating a system of ordinary differential equations corresponding to our model. PS transfer was simulated in non-exchange conditions by considering [PS_A_] = 5 µM, [PS_B_] = 0 µM, [PI4P_A_] = [PI4P_B_] = 0 µM and [O] = 200 nM at t = 0. The concentration of the other forms of the ORD were considered to be equal to 0. PS transfer in exchange conditions was calculated considering that [PI4P_B_] was initially equal to 5 µM. Inversely, PI(4)P transfer was simulated in non-exchange conditions considering [PI4P_B_] = 5 µM, [PI4P_A_] = 0 µM and [PS_A_] = [PS_B_] = 0 µM. In exchange conditions, [PS_A_] value was set at 5 µM. All k_ON-Mb_ and k_OFF-Mb_ rates were set to 10 s^−1^ and 1 s^−1^, respectively. All k_ON-lipid_ and k_OFF-lipid_ rates were set to 10 µM-1. s^−1^ and 1 s^−1^, respectively, unless otherwise specified. The implementation of the kinetic model and the simulations were carried out with the software GEPASI v3.3^61^.

### Cell culture, transfection and drug treatment

HeLa cells were grown in Dulbecco’s modified Eagle medium (DMEM) supplemented with 10% (v/v) fetal bovine serum (Eurobio) at 37°C under 5% CO_2._ Cells were seeded (40,000 cells per condition) in an 8-well coverslip (Ibidi). The next day, the cells were transiently transfected with 125 ng of GFP-C2_Lact_ (Addgene, #22852) plasmid only or additionally with 125 ng of Lyn11-FRB-mCherry plasmid (Addgene, #38004) using Lipofectamine 3000 (Thermo Fisher Scientific) according to the manufacturer’s instructions. To deplete cholesterol in the PM, cells were treated for 24 h with 2.5 µg/mL of U18666A (Sigma) following the transfection step.

### Microscopy and image analysis

One day after transfection, the cells were observed in live conditions using a wide-field microscope (Olympus IX83, 60 ×) or a confocal microscope (Zeiss LSM780, 63 ×). Prior to the observation, the medium was replaced by HEPES-containing DMEM devoid of red phenol. The depletion of cholesterol in the PM was assessed in cells that were only transfected by GFP-C2_Lact_ using the fluorescence sterol sensor mCherry-D4-His_6_. These cells were incubated for 10 min at room temperature with the protein added at 1:500 in DMEM. Then the cells were rinsed twice with HEPES-containing DMEM devoid of red phenol and the cells were immediately observed. To quantify the recruitment of GFP-C2_Lact_ to the PM, line scan analyses of a large set of cells were performed using Fiji ImageJ 2.1.0. A line was manually drawn across each individual cell and the peak intensity at the PM region was normalized with the intensity of the cytosolic region and then plotted for quantification. The localization of the PM was ascertained for each measurement by using Lyn11-FRB-mCherry as an internal reference.

## References

1 Wong, L. H., Gatta, A. T. & Levine, T. P. Lipid transfer proteins: the lipid commute via shuttles, bridges and tubes. Nature reviews. Molecular cell biology 20, 85–101, doi:10.1038/s41580-018-0071-5 (2019).

2 Balla, T., Kim, Y. J., Alvarez-Prats, A. & Pemberton, J. Lipid Dynamics at Contact Sites Between the Endoplasmic Reticulum and Other Organelles. Annual review of cell and developmental biology 35, 85–109, doi:10.1146/annurev-cellbio-100818-125251 (2019).

3 Kumagai, K. & Hanada, K. Structure, functions and regulation of CERT, a lipid-transfer protein for the delivery of ceramide at the ER-Golgi membrane contact sites. FEBS letters 593, 2366–2377, doi:10.1002/1873-3468.13511 (2019).

4 Osawa, T. & Noda, N. N. Atg2: A novel phospholipid transfer protein that mediates de novo autophagosome biogenesis. Protein science : a publication of the Protein Society 28, 1005–1012, doi:10.1002/pro.3623 (2019).

5 Wong, L. H., Čopič, A. & Levine, T. P. Advances on the Transfer of Lipids by Lipid Transfer Proteins. Trends in biochemical sciences 42, 516–530, doi:10.1016/j.tibs.2017.05.001 (2017).

6 Jain, A. & Holthuis, J. C. M. Membrane contact sites, ancient and central hubs of cellular lipid logistics. Biochimica et biophysica acta. Molecular cell research 1864, 1450–1458, doi:10.1016/j.bbamcr.2017.05.017 (2017).

7 Saheki, Y. & De Camilli, P. Endoplasmic Reticulum-Plasma Membrane Contact Sites. Annual review of biochemistry 86, 659–684, doi:10.1146/annurev-biochem-061516-044932 (2017).

8 Grabon, A., Bankaitis, V. A. & McDermott, M. I. The interface between phosphatidylinositol transfer protein function and phosphoinositide signaling in higher eukaryotes. Journal of lipid research 60, 242–268, doi:10.1194/jlr.R089730 (2019).

9 Cockcroft, S. & Raghu, P. Phospholipid transport protein function at organelle contact sites. Current Opinion in Cell Biology 53, 52–60, doi:10.1016/j.ceb.2018.04.011. (2018).

10 Chiapparino, A., Maeda, K., Turei, D., Saez-Rodriguez, J. & Gavin, A. C. The orchestra of lipid-transfer proteins at the crossroads between metabolism and signaling. Progress in lipid research 61, 30–39, doi:10.1016/j.plipres.2015.10.004 (2016).

11 Delfosse, V., Bourguet, W. & Drin, G. Structural and Functional Specialization of OSBP-Related Proteins. Contact 3, 2515256420946627, doi:10.1177/2515256420946627 (2020).

12 Maeda, K. et al. Interactome map uncovers phosphatidylserine transport by oxysterol-binding proteins. Nature 501, 257–261, doi:10.1038/nature12430 (2013).

13 Moser von Filseck, J. et al. INTRACELLULAR TRANSPORT. Phosphatidylserine transport by ORP/Osh proteins is driven by phosphatidylinositol 4-phosphate. Science 349, 432–436, doi:10.1126/science.aab1346 (2015).

14 Chung, J. et al. INTRACELLULAR TRANSPORT. PI4P/phosphatidylserine countertransport at ORP5- and ORP8-mediated ER-plasma membrane contacts. Science 349, 428–432, doi:10.1126/science.aab1370 (2015).

15 Yan, D. et al. OSBP-related protein 8 (ORP8) suppresses ABCA1 expression and cholesterol efflux from macrophages. The Journal of biological chemistry 283, 332–340, doi:10.1074/jbc.M705313200 (2008).

16 Du, X. et al. A role for oxysterol-binding protein-related protein 5 in endosomal cholesterol trafficking. The Journal of cell biology 192, 121–135, doi:10.1083/jcb.201004142 (2011).

17 Ghai, R. et al. ORP5 and ORP8 bind phosphatidylinositol-4, 5-biphosphate (PtdIns(4,5)P 2) and regulate its level at the plasma membrane. Nature communications 8, 757, doi:10.1038/s41467-017-00861-5 (2017).

18 Lee, M. & Fairn, G. D. Both the PH domain and N-terminal region of oxysterol-binding protein related protein 8S are required for localization to PM-ER contact sites. Biochemical and biophysical research communications 496, 1088–1094, doi:10.1016/j.bbrc.2018.01.138 (2018).

19 Sohn, M. et al. PI(4,5)P2 controls plasma membrane PI4P and PS levels via ORP5/8 recruitment to ER-PM contact sites. The Journal of cell biology 217, 1797–1813, doi:10.1083/jcb.201710095 (2018).

20 Di Paolo, G. & De Camilli, P. Phosphoinositides in cell regulation and membrane dynamics. Nature 443, 651–657, doi:10.1038/nature05185 (2006).

21 Behnia, R. & Munro, S. Organelle identity and the signposts for membrane traffic. Nature 438, 597–604, doi:10.1038/nature04397 (2005).

22 Nishimura, T. et al. Osh Proteins Control Nanoscale Lipid Organization Necessary for PI(4,5)P(2) Synthesis. Molecular cell 75, 1043–1057.e1048, doi:10.1016/j.molcel.2019.06.037 (2019).

23 Wenk, M. R. et al. Phosphoinositide profiling in complex lipid mixtures using electrospray ionization mass spectrometry. Nature biotechnology 21, 813–817, doi:10.1038/nbt837 (2003).

24 Klose, C. et al. Flexibility of a eukaryotic lipidome--insights from yeast lipidomics. PLoS One 7, e35063, doi:10.1371/journal.pone.0035063 (2012).

25 Skotland, T. & Sandvig, K. The role of PS 18:0/18:1 in membrane function. Nature communications 10, 2752, doi:10.1038/s41467-019-10711-1 (2019).

26 Symons, J. L. et al. Lipidomic atlas of mammalian cell membranes reveals hierarchical variation induced by culture conditions, subcellular membranes, and cell lineages. Soft Matter 17, 288–297, doi:10.1039/D0SM00404A (2021).

27 Hicks, A. M., DeLong, C. J., Thomas, M. J., Samuel, M. & Cui, Z. Unique molecular signatures of glycerophospholipid species in different rat tissues analyzed by tandem mass spectrometry. Biochim Biophys Acta 1761, 1022–1029, doi:10.1016/j.bbalip.2006.05.010 (2006).

28 Schneiter, R. et al. Electrospray ionization tandem mass spectrometry (ESI-MS/MS) analysis of the lipid molecular species composition of yeast subcellular membranes reveals acyl chain-based sorting/remodeling of distinct molecular species en route to the plasma membrane. The Journal of cell biology 146, 741–754, doi:10.1083/jcb.146.4.741 (1999).

29 Andreyev, A. Y. et al. Subcellular organelle lipidomics in TLR-4-activated macrophages. Journal of lipid research 51, 2785–2797, doi:10.1194/jlr.M008748 (2010).

30 Hirama, T. et al. Membrane curvature induced by proximity of anionic phospholipids can initiate endocytosis. Nature communications 8, 1393, doi:10.1038/s41467-017-01554-9 (2017).

31 Maekawa, M. & Fairn, G. D. Complementary probes reveal that phosphatidylserine is required for the proper transbilayer distribution of cholesterol. Journal of Cell Science 128, 1422, doi:10.1242/jcs.164715 (2015).

32 Bozelli, J. C., Jr. & Epand, R. M. Specificity of Acyl Chain Composition of Phosphatidylinositols. Proteomics 19, e1900138, doi:10.1002/pmic.201900138 (2019).

33 de Saint-Jean, M. et al. Osh4p exchanges sterols for phosphatidylinositol 4-phosphate between lipid bilayers. The Journal of cell biology 195, 965–978, doi:10.1083/jcb.201104062 (2011).

34 Dong, J. et al. Allosteric enhancement of ORP1-mediated cholesterol transport by PI(4,5)P(2)/PI(3,4)P(2). Nature communications 10, 829, doi:10.1038/s41467-019-08791-0 (2019).

35 Wang, H. et al. ORP2 Delivers Cholesterol to the Plasma Membrane in Exchange for Phosphatidylinositol 4, 5-Bisphosphate (PI(4,5)P(2)). Molecular cell 73, 458–473.e457, doi:10.1016/j.molcel.2018.11.014 (2019).

36 D’Ambrosio, J. M. et al. Osh6 requires Ist2 for localization to ER-PM contacts and efficient phosphatidylserine transport in budding yeast. J Cell Sci 133, doi:10.1242/jcs.243733 (2020).

37 Zhang, Y. et al. High-Throughput Lipidomic and Transcriptomic Analysis To Compare SP2/0, CHO, and HEK-293 Mammalian Cell Lines. Anal Chem 89, 1477–1485, doi:10.1021/acs.analchem.6b02984 (2017).

38 Llorente, A. et al. Molecular lipidomics of exosomes released by PC-3 prostate cancer cells. Biochim Biophys Acta 1831, 1302–1309, doi:10.1016/j.bbalip.2013.04.011 (2013).

39 Kavaliauskiene, S. et al. Cell density-induced changes in lipid composition and intracellular trafficking. Cellular and molecular life sciences : CMLS 71, 1097–1116, doi:10.1007/s00018-013-1441-y (2014).

40 Dickson, E. J. et al. Dynamic formation of ER-PM junctions presents a lipid phosphatase to regulate phosphoinositides. The Journal of cell biology 213, 33–48, doi:10.1083/jcb.201508106 (2016).

41 Traynor-Kaplan, A. et al. Fatty-acyl chain profiles of cellular phosphoinositides. Biochim Biophys Acta Mol Cell Biol Lipids 1862, 513–522, doi:10.1016/j.bbalip.2017.02.002 (2017).

42 Lemmon, M. A., Ferguson, K. M., O’Brien, R., Sigler, P. B. & Schlessinger, J. Specific and high-affinity binding of inositol phosphates to an isolated pleckstrin homology domain. Proc Natl Acad Sci U S A 92, 10472–10476, doi:10.1073/pnas.92.23.10472 (1995).

43 Ferguson, K. M., Lemmon, M. A., Schlessinger, J. & Sigler, P. B. Structure of the high affinity complex of inositol trisphosphate with a phospholipase C pleckstrin homology domain. Cell 83, 1037–1046, doi:10.1016/0092-8674(95)90219-8 (1995).

44 Lipp, N.-F. et al. An electrostatic switching mechanism to control the lipid transfer activity of Osh6p. Nature communications 10, 3926, doi:10.1038/s41467-019-11780-y (2019).

45 Drin, G. Topological regulation of lipid balance in cells. Annual review of biochemistry 83, 51–77, doi:10.1146/annurev-biochem-060713-035307 (2014).

46 Amblard, I. et al. Bidirectional transfer of homeoprotein EN2 across the plasma membrane requires PIP2. J Cell Sci 133, doi:10.1242/jcs.244327 (2020).

47 Underwood, K. W., Jacobs, N. L., Howley, A. & Liscum, L. Evidence for a cholesterol transport pathway from lysosomes to endoplasmic reticulum that is independent of the plasma membrane. The Journal of biological chemistry 273, 4266–4274, doi:10.1074/jbc.273.7.4266 (1998).

48 Huuskonen, J. et al. Acyl chain and headgroup specificity of human plasma phospholipid transfer protein. Biochimica et Biophysica Acta (BBA) - Lipids and Lipid Metabolism 1303, 207–214, doi:10.1016/0005-2760(96)00103-8 (1996).

49 Backman, A. P. E. et al. Glucosylceramide acyl chain length is sensed by the glycolipid transfer protein. PLoS One 13, e0209230, doi:10.1371/journal.pone.0209230 (2018).

50 Kudo, N. et al. Structural basis for specific lipid recognition by CERT responsible for nonvesicular trafficking of ceramide. Proc Natl Acad Sci U S A 105, 488–493, doi:10.1073/pnas.0709191105 (2008).

51 Kumagai, K. et al. CERT mediates intermembrane transfer of various molecular species of ceramides. The Journal of biological chemistry 280, 6488–6495, doi:10.1074/jbc.M409290200 (2005).

52 Silvius, J. R. & Leventis, R. Spontaneous interbilayer transfer of phospholipids: dependence on acyl chain composition. Biochemistry 32, 13318–13326, doi:10.1021/bi00211a045 (1993).

53 Singh, R. P., Brooks, B. R. & Klauda, J. B. Binding and release of cholesterol in the Osh4 protein of yeast. Proteins 75, 468–477, doi:10.1002/prot.22263 (2009).

54 Canagarajah, B. J., Hummer, G., Prinz, W. A. & Hurley, J. H. Dynamics of cholesterol exchange in the oxysterol binding protein family. Journal of molecular biology 378, 737–748, doi:10.1016/j.jmb.2008.01.075 (2008).

55 Manni, M. M. et al. Acyl chain asymmetry and polyunsaturation of brain phospholipids facilitate membrane vesiculation without leakage. Elife 7, doi:10.7554/eLife.34394 (2018).

56 Mesmin, B. et al. A four-step cycle driven by PI(4)P hydrolysis directs sterol/PI(4)P exchange by the ER-Golgi tether OSBP. Cell 155, 830–843, doi:10.1016/j.cell.2013.09.056 (2013).

57 Zhong, W. et al. ORP4L Extracts and Presents PIP_2_ from Plasma Membrane for PLCβ3 Catalysis: Targeting It Eradicates Leukemia Stem Cells. Cell Reports 26, 2166–2177.e2169, doi:10.1016/j.celrep.2019.01.082 (2019).

58 Moser von Filseck, J., Vanni, S., Mesmin, B., Antonny, B. & Drin, G. A phosphatidylinositol-4-phosphate powered exchange mechanism to create a lipid gradient between membranes. Nature communications 6, 6671, doi:10.1038/ncomms7671 (2015).

59 Morales, J., Sobol, M., Rodriguez-Zapata, L. C., Hozak, P. & Castano, E. Aromatic amino acids and their relevance in the specificity of the PH domain. Journal of Molecular Recognition 30, e2649, doi:10.1002/jmr.2649 (2017).

60 Alexandrov, A. I., Mileni, M., Chien, E. Y., Hanson, M. A. & Stevens, R. C. Microscale fluorescent thermal stability assay for membrane proteins. Structure 16, 351–359, doi:10.1016/j.str.2008.02.004 (2008).

61 Mendes, P. GEPASI: a software package for modelling the dynamics, steady states and control of biochemical and other systems. Comput Appl Biosci 9, 563–571 (1993).

